# IFN-associated B cell hyperactivity is highly enriched in SLE patients harboring Sm/RNP antibodies

**DOI:** 10.1101/2024.11.18.624119

**Authors:** H.J. van Dooren, Y. Atisha-Fregoso, A.L. Dorjée, T.W.J. Huizinga, M. Mackay, C. Aranow, R.E.M. Toes, B. Diamond, J. Suurmond

## Abstract

Systemic Lupus Erythematosus (SLE) is an autoimmune disease characterized by an array of autoantibodies, in particular anti-nuclear antibodies (ANA). The disease is also hallmarked by an expansion of plasmablasts (PB) in peripheral blood. How these relate to autoantibody production is not clear. Here, we aimed to understand B cell alterations in SLE and their relationship to immunoglobulin levels and autoantibody production.

We demonstrate that a subgroup of SLE patients is characterized by a high frequency of PB relative to memory B cells (high PB/M). Patients with this phenotype more frequently had high disease activity. Despite low overall frequencies of memory B cells, these patients exhibited an increased activation in the switched CD27+ memory compartment and a strong IFN signature in PB. Repertoire analysis revealed a highly polyclonal expansion and enrichment for IgG1 expressing PB in patients with a high PB/M ratio which was reflected in increased serum IgG levels. Importantly, the hyperactive B cell phenotype was highly enriched in patients harboring Sm/RNP autoantibodies (OR: 9.17 (2.97-26.0)). In summary, we show for the first time a direct relationship between IFN and PB expansion in a subgroup of SLE patients, highly enriched in those harboring Sm/RNP antibodies. These results provide insight into the pathways leading to B cell hyperactivity and autoantibody production which may guide the tailoring of B cell- and IFN-targeted therapies.

**Graphical abstract:** 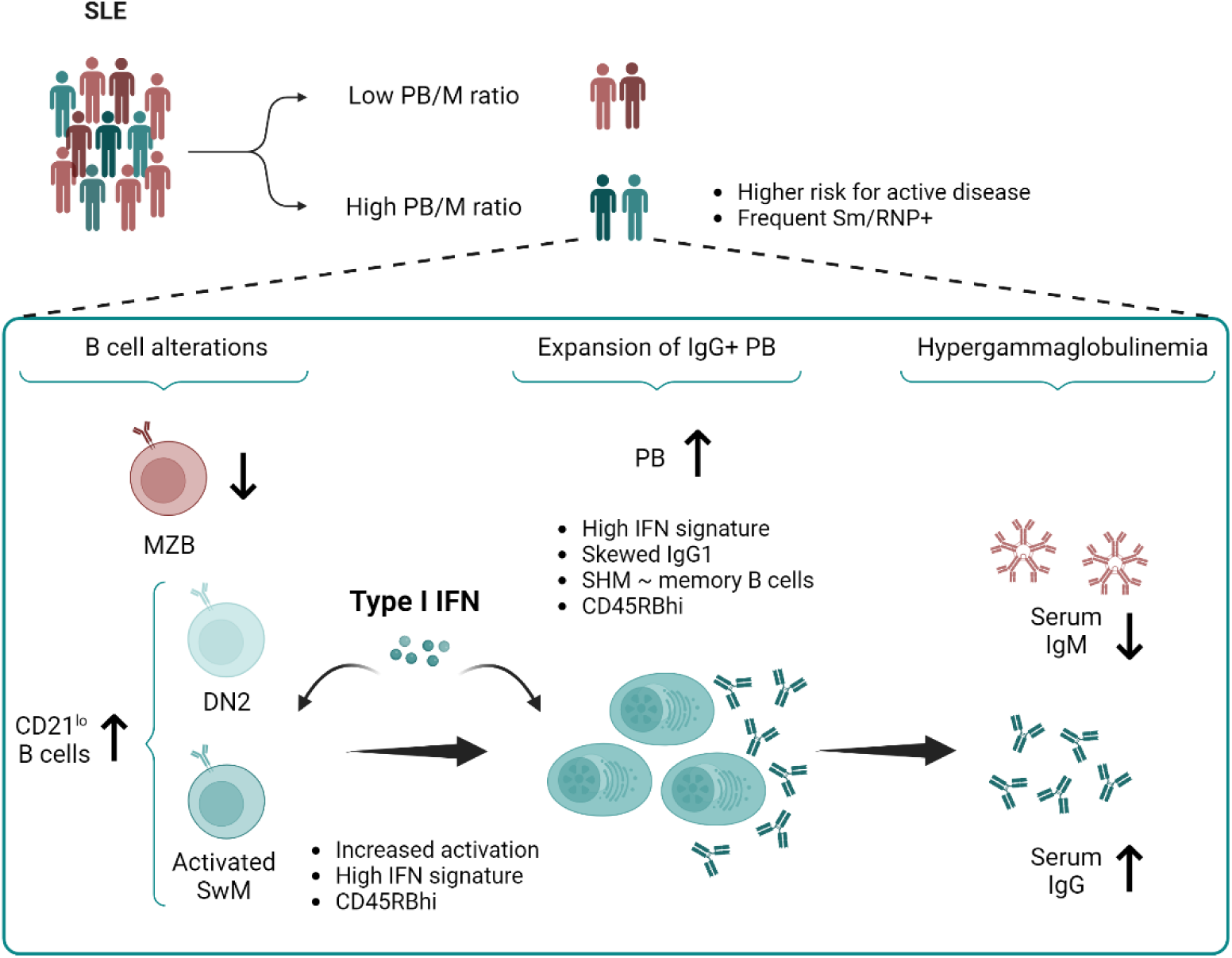

## INTRODUCTION

Systemic Lupus Erythematosus (SLE) is a systemic autoimmune disease characterized by autoantibodies and chronic inflammation. It is a heterogeneous disease that can affect multiple organs and clinical activity can fluctuate considerably (1). The presence of antinuclear antibodies (ANA) is a hallmark of SLE. ANA consist of antibodies that can target a large variety of nuclear antigens (2). The most well-known antigens are double-stranded DNA (dsDNA), and RNA binding proteins (RBPs) that can be further divided into Sm/RNP complex and SS-A/SS-B antigens. The presence of anti-dsDNA and anti-Sm are the most specific for SLE and are part of classification criteria (3). Other ANA reactivities can be present in other autoimmune diseases (2).

ANA are produced by plasmablasts/plasma cells (PB/PC), which can arise through various B cell differentiation pathways (4). The anti-chromatin and anti-RBP antibodies are often considered to arise through distinct pathways, anti-dsDNA titers fluctuate and are associated most strongly with disease flares and lupus nephritis, whereas most anti-RBPs are more stable (5, 6). Anti-dsDNA levels strikingly decreased following CD19 CAR T cell treatment while long-lived vaccination responses remained intact (7), suggesting an important role for extrafollicular B cell activation, at least for some of the ANAs. However, the exact pathways that are involved in the production of ANAs are not known. In particular, it is not clear whether and how global B cell alterations associate with antigenic specificity.

The most well-known feature of the B cell population in SLE is the presence of a high number of circulating PB, in particular of the IgG isotype (8–15). Higher numbers of circulating PB are usually found in a subgroup of patients, and a high PB frequency or a high transcriptomic PB signature is associated with active disease or flares (11, 12, 14). The mechanism for expansion of circulating PB is unclear. Such understanding is hampered by the sensitivity of PB to cryopreservation, making the analyses of PB more demanding as it requires the availability of fresh blood. Inducing B cell hyperactivity through single gene variants or knockouts in mouse models often leads to PC expansion and hypergammaglobulinemia, while also inducing specific autoantibodies (16, 17). Genetic variants leading to B cell hyperactivity have been reported for monogenic SLE in humans as well (18, 19). These gene variants often affect B cell signaling through the BCR or TLR, and as such, are expected to increase the response not only to self-antigens, explaining the heightened serum IgG levels. In line with this, altered function of B cell signaling as seen in SLE can augment B cell responses to both self- and foreign antigens (16–18). B cell hyperactivity is causally involved in the loss of B cell tolerance and SLE. However, how PB expansion relates to specific ANA in human SLE is not clear.

Previously, we identified that patients with a high frequency of ANA+ PB usually have a low frequency of ANA+ CD27+ memory B cells (20). This phenotype was particularly present in a subgroup of SLE patients, but we did not yet consider the global B cell alterations and the relationship to global PB expansion in these patients. Besides PB expansion, virtually all other B cell subsets can be altered in frequency in SLE, including double-negative 2 (DN2) cells, age- or autoimmunity-associated B cells (ABCs), activated naïve B cells, MZ-like B cells, and transitional B cells (21–24). The mechanisms underlying alterations in B cell subsets and the relationship of each to PB expansion or differentiation is unclear. In addition, it is unknown which of the alterations in the distribution of the B cell compartment lead to increased antibody production, both for total serum immunoglobulins as well as for specific autoantibodies.

In this study we aimed to characterize B cell alterations in SLE and their relationship to immunoglobulin and autoantibody production, with the ultimate goal to identify features andmechanisms underlying B cell hyperactivity and autoantibody production. Our results reveal a novel distribution of the B cell compartment that associates with B cell hyperactivity, a phenotype that is particularly prominent in patients with active disease. This phenotype is characterized by a high PB to memory B cell ratio, increased activation in the switched memory compartment, and an IFN- associated polyclonal expansion of the PB compartment. A high PB to memory B cell ratio marks patients with increased ratios of IgG:IgM and Sm/RNP autoantibodies. Thus increased differentiation of switched B cells towards PB likely contributes to increased antibody production in SLE.

## RESULTS

### A high PB to memory B cell ratio characterizes a subgroup in SLE with high disease activity

Several studies report increased numbers of circulating PB in SLE. In these studies, the relative frequency of PB out of total B cells (%PB) as well as the absolute PB count are frequently found to be increased (11, 12, 14). Using principal component analysis we previously found a distinct SLE patient group with a high frequency of ANA+ PB relative to the frequency of ANA+ CD27+ memory B cells (Bmem) (20). We now investigated whether a similar PB to memory ratio was also observed in the total B cell compartment. Consistent with literature, we observed a trend for a higher frequency of PB, particularly visible in a subset of SLE patients (Figure 1A,B). As we and others observed before, SLE was also characterized by a significantly reduced frequency of CD27+ Bmem (Figure 1C) (16, 20). Patients with a high frequency of PB tended to have a lower frequency of CD27+ Bmem (Figure 1D). The ratio of PB to CD27+ Bmem (PB/M ratio) in SLE patients was significantly increased compared to healthy donors (Figure 1E). As a subset of patients exhibited this phenotype, we divided SLE patients into subgroups with a low and high PB/M ratio, with a cutoff based on the healthy donors (Q3 + 1.5*IQR). The PB/M ratio was highly stable over time as repeated measurements approximately 1 year apart available in the US cohort (n=24) revealed a strong correlation (Figure 1F). These findings were replicated in an independent cohort in Europe (Figure S1).

**Figure 1:**
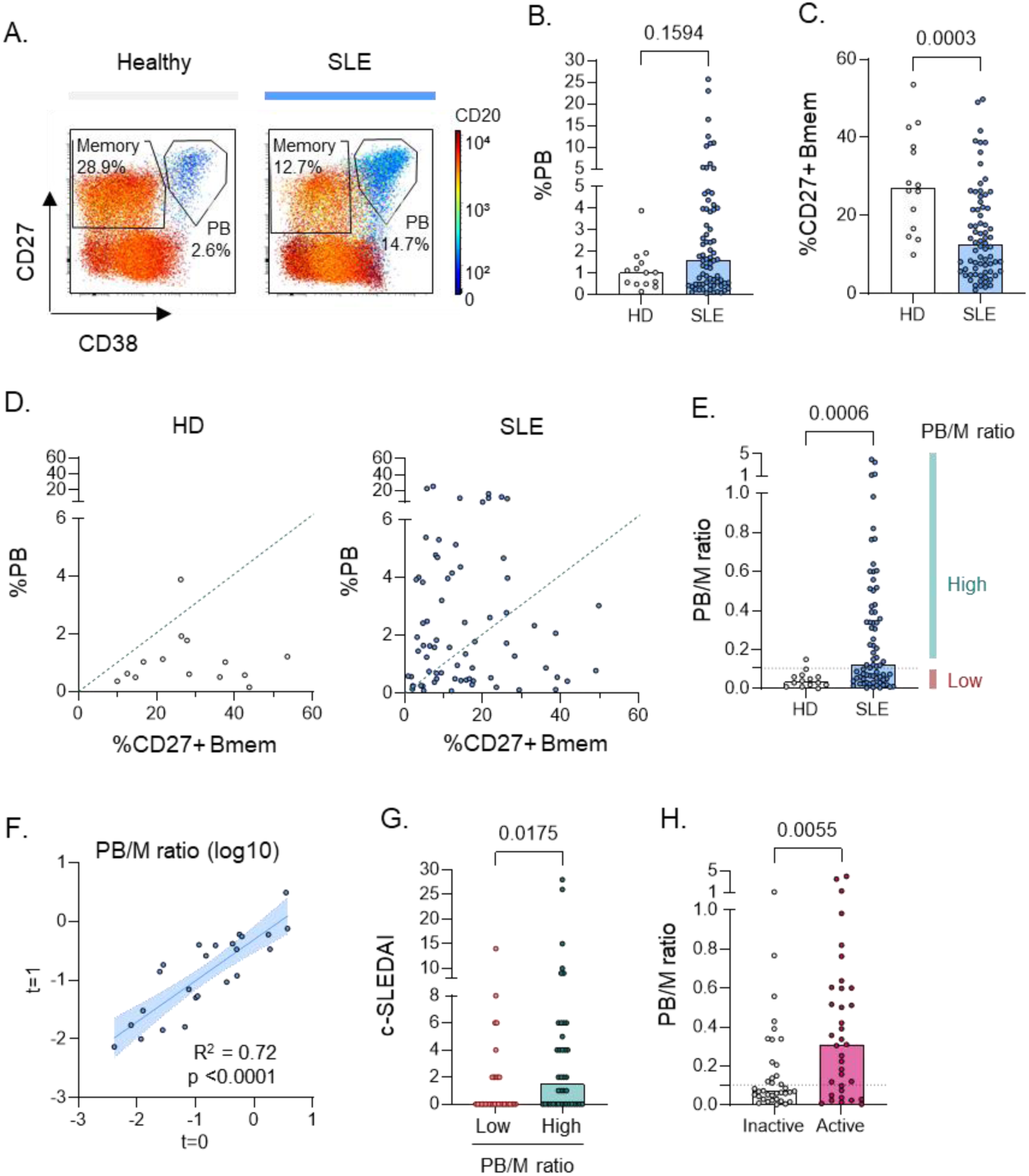
A high PB to memory B cell ratio characterizes a subgroup in SLE with high disease activity. B cell phenotypes of SLE patients (n=72) and healthy donors (n=14) were analysed using flow cytometry. A) Representative examples of PB and Bmem gating. B-C) Percent PB and CD27+ Bmem among total B cells. D) Scatterplot of %PB and %CD27+ Bmem with the dashed line representing a PB/M ratio of 0.105. E) Ratio of PB over CD27+ Bmem (PB/M). F) Correlation of PB/M ratio over time in SLE patients whose B cell phenotype was measured twice (n=24). G) c-SLEDAI in SLE patients with a low versus high PB/M ratio. H) PB/M ratio in patients with clinically inactive versus active (c-SLEDAI>0) disease. Each dot indicates an individual, and the bars represent the median. P values were calculated using Mann- Whitney test (B,C,E,G,H), or linear regression (F).

Analysis of patient characteristics of the “Low PB/M” and “High PB/M” groups in the US cohort (Table S1,S2) revealed trends for a higher frequency of male/female (OR: 6.576; 95%CI: 1.034-76.14) and self-reported African-American race (OR: 2.778; 95% CI: 1.104-6.827) in the high PB/M group, though these were not statistically significant and not reproduced in the replication cohort. The only significant difference was observed for Asian race, which was more abundant in the low PB/M group, though this was based on small numbers of patients in the US cohort. We next addressed whether patients with a high PB/M ratio display a different clinical profile. Patients with a high PB/M ratio had a higher SLEDAI-2k disease activity score and more frequently moderate to high disease activity (Figure 1G,H, Table S3). Clinical activity was observed across different disease manifestations, though not statistically significant for most symptoms (Table S4). The exception was hematological disease activity, in which patients with a high PB/M ratio more frequently had leukopenia and lymphopenia, but not thrombocytopenia (Figure S2A-D). Patients with a high PB/M ratio also displayed decreased levels of complement C4, but not C3 (Figure S2E,F). No difference in disease activity was observed when analyzing SLE patients by frequency or absolute numbers of PB (Figure S2G,H). No clear differences with regards to medication were observed (Table S1,S2).

Though the PB/M ratio was relatively stable over time (Figure 1E), we also analysed whether a change in PB phenotype associated with a change in disease activity. No clear correlation between a change in disease activity (delta SLEDAI-2k) and change in PB parameters (delta PB/M ratio or delta %PB) was observed (Figure S2I-L), possibly due to low patient numbers (n=24).

Overall, these results suggest a stable phenotype of high PB/M ratio that is associated with a higher risk for clinically active disease.

A high frequency of PB relative to memory B cells (PB/M) is present in a subgroup of SLE patients with a higher disease activity. This more strikingly differentiates between SLE patients and healthy subjects than the frequency or number of PB.

### Increased activation within switched B cell compartments

To obtain further understanding of the alterations of B cell populations in SLE, we performed detailed spectral flow cytometry. Based on unsupervised clustering with 8 B cell markers (CD19, CD20, CD27, CD38, CD21, CD24, IgD, IgM), we identified 13 B cell subsets (Figure 2A,B, Figure S3A). Clusters were assigned to B cell populations according to published definitions (25). Using the same cutoff (0.105) for the PB/M ratio to subset SLE patients (Figure 2C), we first analysed the distribution of subsets within the CD27+ memory compartment (Figure 2D,E), revealing a strong decrease in resting unswitched memory B cells (CD27+CD38^lo^CD21+CD24+IgD+IgM+) in SLE patients with a high PB/M ratio. These cells have also been referred to as marginal zone-like cells (16, 25, 26). Despite the overall reduction in CD27+ memory B cells, an increase in the activated switched memory compartment was observed. These cells were characterized by increased expression of CD19, CD20, intermediate CD38, and low CD21 expression compared to resting switched memory B cells (Figure 2F). Within the CD27- B cell subsets a significant decrease in resting naïve B cells was observed in patients with a high PB/M ratio (Figure S3B). Besides a relative increase in activation within the SwM compartment, Pre-PB, cells with an intermediate phenotype between Act SwM and PB, were also increased in patients with a high PB/M ratio (Figure S3C,D).

**Figure 2:**
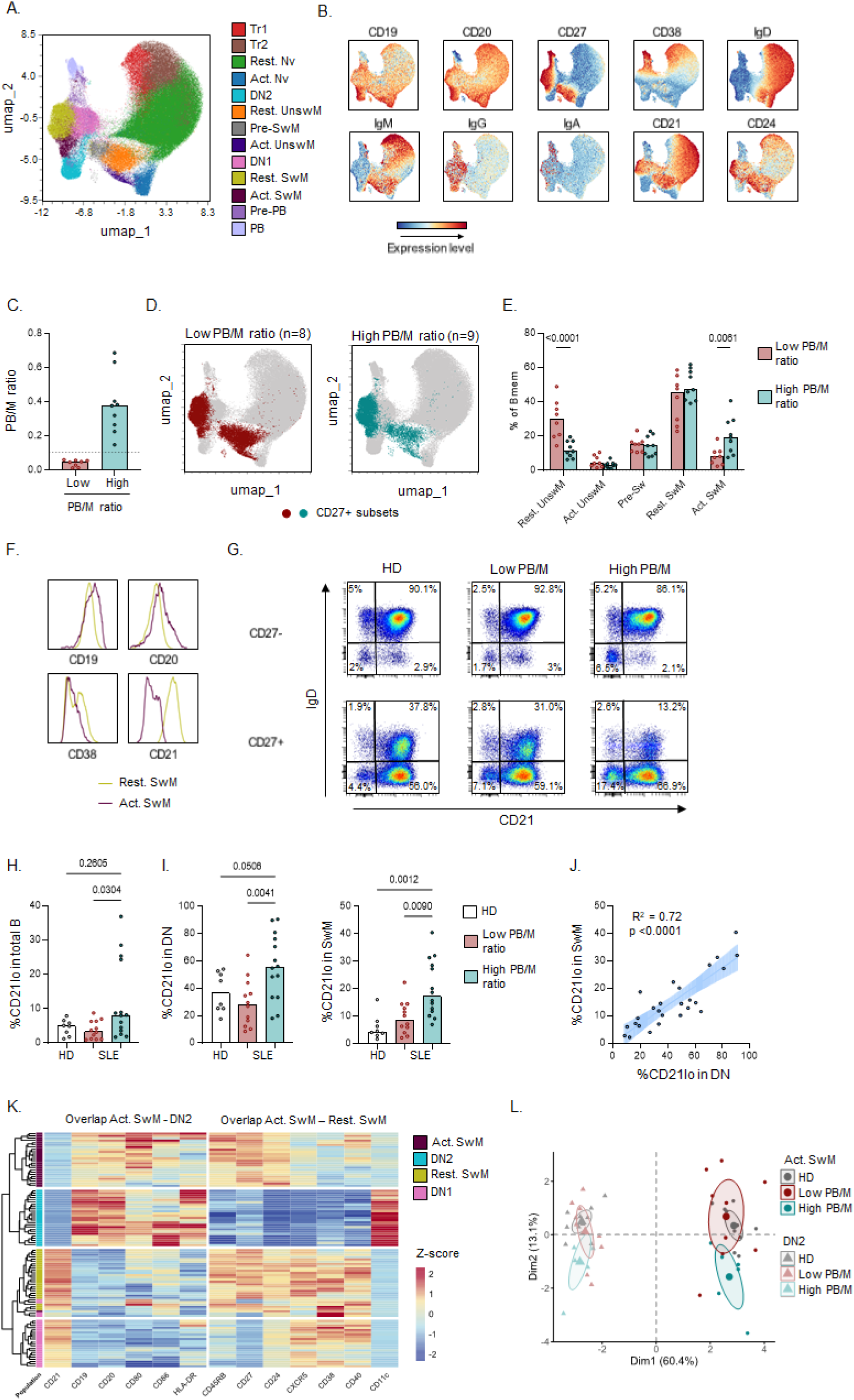
Increased activation within switched B cell compartments. High dimensional spectral flow cytometry was used for detailed B cell phenotyping within fresh PBMCs (A-F) from SLE patients (SLE; n=17) and a second cohort of frozen PBMCs (K-L) from SLE patients (n=15) and healthy donors (HD; n=10). Data from both experiments was pooled in H-L, after calculating the average for 5 samples present in both. Additional data on both cohorts is provided in Figure S3 and S4. A) Live B cells were clustered using FlowSOM and visualized in UMAP. B) Normalized expression levels of B cell markers in UMAP plot. C) PB/M ratio in this dataset. D) Projection of CD27+ Bmem cells on the UMAP plot. E) %clusters within the CD27+ Bmem cells in SLE patients with a low versus high PB/M ratio. F) Histograms of key markers defining the Act SwM cluster. G) IgD and CD21 expression in CD27- and CD27+ B cells. H) %CD21^lo^ within total B cells. I) %CD21lo within switched B cells (DN and SwM subsets). J) Correlation of %CD21lo within DN and SwM B cells in SLE patients. K) Clustered heatmap of expression level of activation markers in 4 switched B cell subsets in the second dataset. Expression level is represented as a z-score of the scaled median fluorescence intensity per subset per sample. L) Principal component analysis of all B cell markers from panel K. The percentage indicated on the axis is the % of variance explained by that principal component. The ellipse depicts the 95% confidence interval. Each dot indicates an individual, and the bars represent the median. P values were calculated using Mann- Whitney test (C), Kruskal-Wallis with FDR posthoc test (H,I), simple linear regression (J) or Two-way ANOVA with FDR posthoc test (E).

To further confirm the findings described above, we next generated an additional dataset using a more detailed spectral flow cytometry panel including several activation markers on frozen PBMCs (16). Despite the relative reduction of PB in frozen cells (Figure S4A,B), the main differences in B cell subset distribution in SLE compared to HD were confirmed (Figure S4C-H). Though no significant differences were observed in the percentages of DN2 out of total B cells in both datasets, when combining both datasets and focusing on the CD21^lo^ cells within each subset, both CD27- and CD27+ cells displayed increased percentages of CD21^lo^ cells, most prominently in the switched compartments (DN and SwM respectively) (Figure 2G-I). These subsets represent DN2 and activated SwM B cells, respectively. The frequency of CD21^lo^ cells within these subsets was highly correlated (Figure 2J), suggesting that activation across switched B cells is common in SLE patients, in particular those with a high PB/M ratio. Since CD21^lo^ cells have been implicated in SLE before, we analysed the expression level of activation markers in the different switched B cell clusters with low CD21 expression and their CD21+ counterparts (DN2 vs DN1, and Activated SwM vs Resting SwM). Hierarchical clustering revealed that Act SwM cells have similarities with DN2 cells (decreased CD21, increased CD19, CD20, CD80, CD86, HLA-DR) confirming their activated status, even though sharing marker expression with conventional CD27+ memory B cells (increased CD45RB, CD27, CD24, CXCR5, CD38, CD40 and low CD11c) (Figure 2K). High CD45RB expression has been previously linked to a germinal center origin (27), and CD45RB expression in PB was high and comparable to that of CD27+ memory B cell subsets in both patient groups (Figure S5A). Using principal components analysis of activation markers, the phenotype of Act SwM in patients with a high PB/M ratio was distinct from that of patients with a low PB/M ratio (Figure 2L), and was characterized by decreased expression of CD21 and increased expression of CD38, CD86, CD11c, and a trend towards increased expression of CD40 and CD80 (Figure S5B,C). Together, these data indicate that both the frequency and degree of activation of CD21^lo^ switched memory B cells are increased in patients with a high PB/M ratio. Though DN2 cells displayed a less divergent phenotype, CD86 and CD11c were also expressed at significantly higher levels in DN2 cells from patients with a high PB/M ratio (Figure S5B,C). Using trajectory analysis starting at Tr1 cells, the DN2, Act SwM and Pre-PB populations were those closest to PB (Figure S3E-G). Thus, a high PB/M ratio is related to B cell hyperactivity in switched B cells with skewing of B cells towards the most activated and differentiated subsets.

### Increased proliferation and IFN signature underly PB expansion in patients with a high PB/M ratio

To identify the mechanism underlying PB differentiation in these patients, we performed scRNAseq of FACS sorted PBs (CD19+CD27++CD38++) from SLE patients with a low (n=4) versus high (n=5) PB/M ratio, in 2 independent experiments. After QC filtering based on number of genes detected per cell, percentage of mitochondrial genes, contaminating cells (non-PB) and doublets (see Methods), there were a total of 1290 and 1197 PB analysed in the two experiments respectively. Using unsupervised clustering and visualization, the clusters and differentially expressed genes largely overlapped between the datasets (data not shown), and were correlated to the patient B cell phenotype in both datasets. We therefore proceeded with an integrated analysis of both datasets. Using unsupervised clustering, 2 distinct clusters could be identified (Figure 3A). Genes upregulated in cluster 1 were associated with proliferation/cell division (Figure 3B,C, Figure S6A,B), and this cluster was increased in frequency in patients with a high PB/M ratio (Figure 3D,E). Importantly, besides this increased abundance in cluster 1, covarying neighborhood analysis revealed that cells from patients in each group were closely related to cells from their own group despite being represented in one cluster (Figure 3F,G). Differential gene expression revealed increased expression of IFN-stimulated genes in PBs from patients with a high PB/M ratio (Figure 3H,I, Figure S6C). Whereas IFN-stimulated genes between type I IFN and IFN-gamma largely overlap, enrichment in transcription factor binding sites for IRF7 and ISRE (Figure S6D), as well as expression levels of IFN receptor units (Figure S6E) suggest this signature is most likely related to type I IFN.

**Figure 3:**
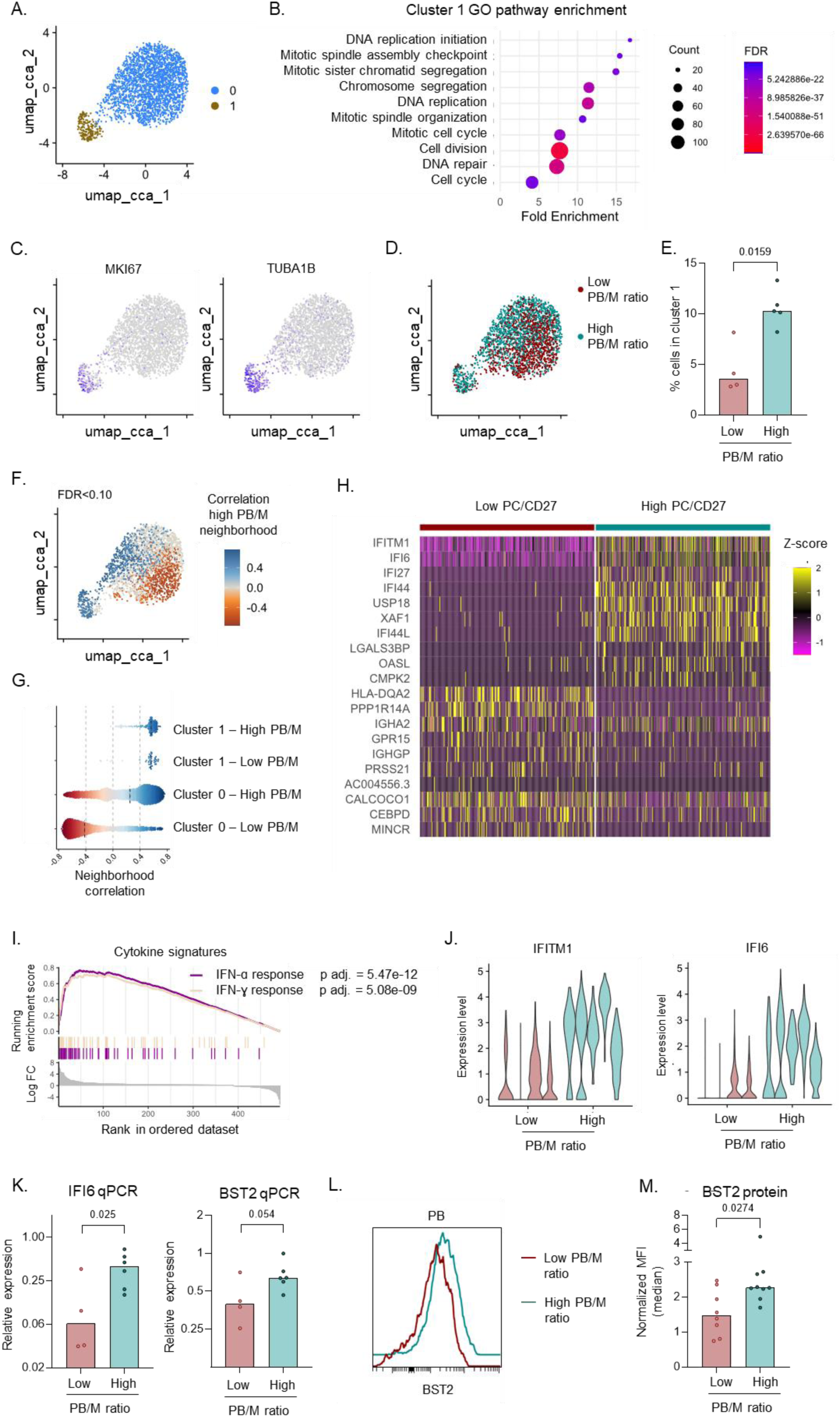
Increased proliferation and IFN signature underly PB expansion in patients with a high PB/M ratio. scRNA-seq was performed on sorted PB from 9 SLE patients, in 2 independent experiments. A) UMAP using canonical correlation analysis for integration of both datasets with 2 clusters identified using unsupervised clustering. B) Pathway enrichment in genes upregulated in cluster 1. C) Expression level of MKI67 and TUBA1B on UMAP. D) Distribution of cells in patients with a low versus high PB/M ratio. E) % cells in cluster 1 between the two patient groups. F,G) Results of association test for a high PB/M ratio using CNA (CNA global P = 0.030): each cell with an FDR> 0.010 is colored according to its neighborhood coefficient, with blue indicating a positive correlation and red indicating a negative correlation (F), and each cell is split per patient group and per cluster (G). H) Heatmap of top 10 differentially expressed genes (adjusted p value <5*10e-5, highest fold change) between patients with a low versus high PB/M ratio. 250 cells from each group are displayed. I) GSEA for cytokine signatures using differentially expressed genes (adjusted p value <0.05; ordered by log fold change) between the two patient groups. J) Expression level of IFI6 and IFITM1 in patients with a low versus high PB/M ratio. Each patient is displayed as individual “violin”. K) RNA expression of IFI6 and BST2 in sorted PB, determined by qPCR. Relative expression was calculated based on two housekeeping genes. L,M) BST2 protein expression on PB was determined using spectral flow cytometry as described in Figure 2. L) Representative example. M) Normalized expression of BST2 in patients with a low versus high PB/M ratio. Each dot in the bargraphs indicates an individual, and the bars represent the median. P values and FDR values were calculated using DAVID functional annotation (B), Mann-Whitney test (E,M), CNA (F), GSEA (I), or student’s T-test on log-normalized expression (K).

The increased IFN signature was observed in each individual patient from the high PB/M group (Figure 3J) and was confirmed in additional patients by qPCR (Figure 3K). Furthermore, increased protein expression of BST2, an IFN-regulated cell surface marker, was confirmed on PBs by spectral flow cytometry (Figure 3L,M). Within the PB/M high patient group, high BST2 expression was shared among all CD21^lo^ B cell subsets (activated naïve, DN2, activated unswitched and activated switched memory) as well as pre-PB and PB (Figure S7A). However, high expression of BST2 in CD21lo subsets was also found in healthy donors and SLE patients with a low PB/M ratio. No significant differences in BST2 expression were found between groups for CD21lo subsets (Figure S7B). In contrast, Pre-PB and PB showed significantly increased BST2 expression in SLE patients with a high PB/M ratio (Figure S7C). This suggests that IFN is highly associated with accelerated differentiation of PBs in patients with a high PB/M ratio.

### IgG1 skewing and highly polyclonal BCR repertoire in PB

We next analysed the BCR repertoire obtained through scRNAseq to obtain insight into the cellular origin of PB in patients with a high PB/M ratio. First, we discovered that patients with a high PB/M ratio displayed a skewing in the isotype distribution, with decreased IgA and increased IgG abundance (Figure 4A). In addition, we observed increased IgG1 and decreased IgG2 within the IgG+ cells in this group (Figure 4B). Decreased IgA and increased IgG abundance in PB was confirmed in spectral flow cytometry (Figure 4C,D). Interestingly, IgG1+ cells displayed the most prominent IFN signature with highest expression of IFN-stimulated genes IFI6 and IFITM1 (Figure 4E). IgG1 cells had a similar expression level of IFNAR1/2 as other subclasses (Figure S8A), suggesting the increased IFN signature is due to their local activation/differentiation by IFN.

**Figure 4:**
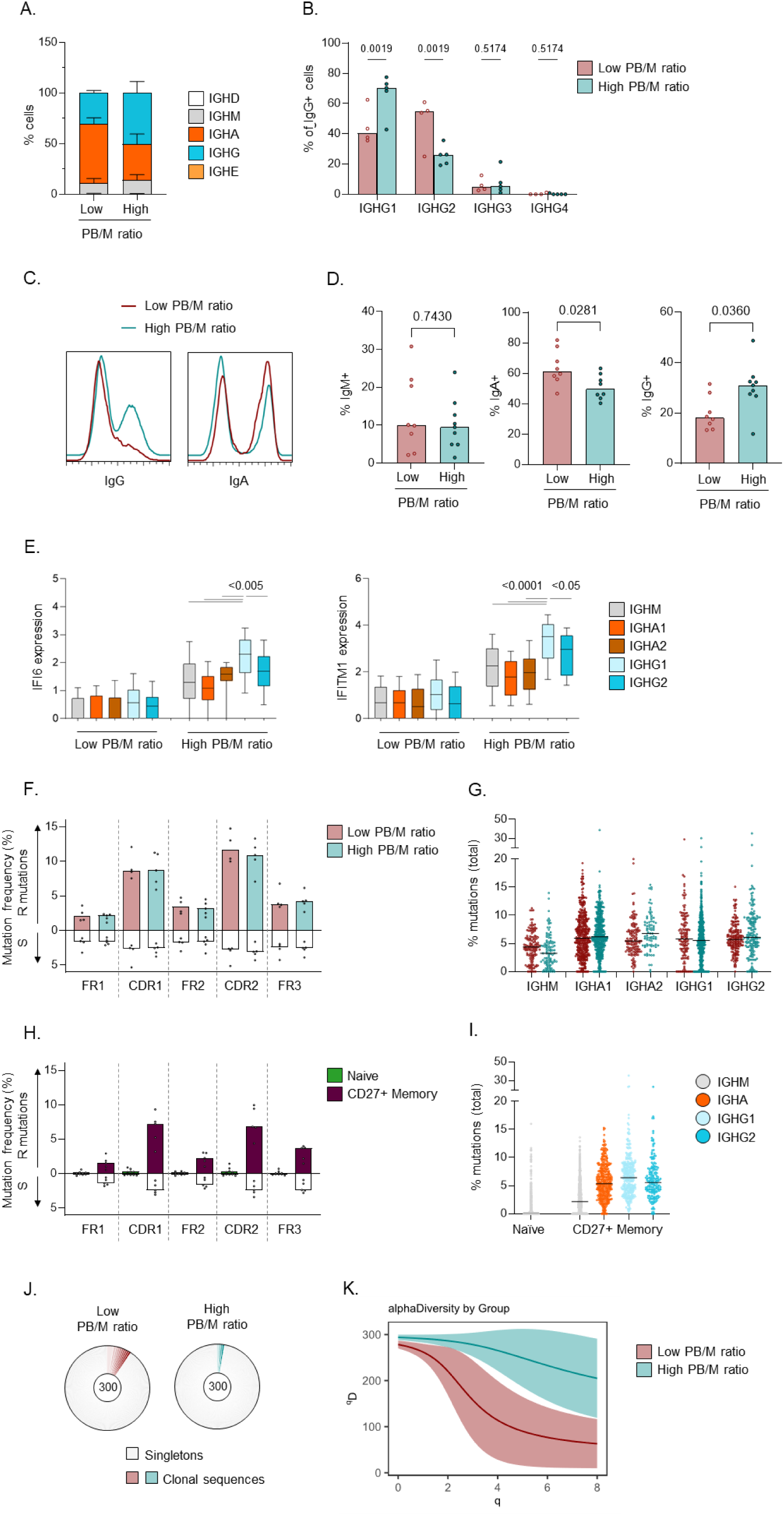
IgG1 skewing and highly polyclonal BCR repertoire in PB. BCR repertoire was determined using results from scRNA-seq (Figure 3) from 9 SLE patients, in 2 independent experiments. A) Frequency of IGH isotypes in PB from patients with a low versus high PB/M ratio. B) IGHG subclass distribution within IgG+ cells in both patient groups. C,D) Expression of IgM, IgA, and IgG in PB was determined using spectral flow cytometry as described in Figure 2. C) Representative examples for IgA and IgG. D) Percent of each isotype in PB from patients with a low versus high PB/M ratio. E) scRNAseq expression level of IFI6 and IFITM1 in cells from each IGH isotype/subclass, separated by patient group. IGHD, IGHG3, and IGHG4 cells were not analysed due to low number of cells (panel A). F,H) % replacement (R) and silent (S) mutations, per region, in PB from patients with a low versus high PB/M ratio (F) and in naïve and CD27+ memory B cells from SLE patients (H). G,I) Total % mutations in PB by Ig subclass in PB (G) and in naïve and CD27+ memory B cells from SLE patients (I). J) Expanded clones and singletons in PB from both patient groups. scRNAseq PB data was downsampled to 100 cells/patient and the total of 3 patients is shown. K) Smoothed alpha diversity (ɑD) curve over a range of diversity orders (q) in data from panel J. Data in are shown as mean + SEM (A), as median across individual patients (B,D,F,H), as boxplot with interquartile range (E), as individual cells with the median across all cells (G,I), or as mean with the 95% confidence interval K). P values were calculated using Two-way ANOVA with FDR posthoc test (B,F-G), Mann-Whitney test (D), or One-way ANOVA with FDR posthoc test (E). No significant differences were found between patients in F,G.

No obvious skewing of V gene usage was observed (Figure S8B). No difference in somatic hypermutation (SHM) was observed between the groups, whether analysed by location in the V-gene or by isotype/subclass (Figure 4F,G). The frequencies of SHM were similar to those observed for memory B cells (Figure 4H,I). Together with the high R/S ratio in CDR regions this suggests that PB in both groups originate from cells that underwent selection processes, most likely in the germinal center. Analysis of clonality revealed increased diversity in patients with a high PB/M ratio compared to those with a low PB/M ratio (Figure 4J,K).

Together, these analyses indicate that PB in patients with a high PB/M ratio arise through IFN-driven polyclonal differentiation and/or expansion of IgG1+ PB likely derived from post-germinal center cells.

### B cell hyperactivity results in hypergammaglobulinemia

Hypergammaglobulinemia is often regarded as a sign of B cell hyperactivity. We observed decreased levels of IgM and increased levels of IgA and IgG in SLE compared to healthy controls, resulting in a high ratio of IgG:IgM in SLE patients (Figure 5A-D). These features were most pronounced in SLE patients with a high PB/M ratio (Figure 5E-H). These findings were replicated in an independent second cohort (Figure S9A-H).

**Figure 5:**
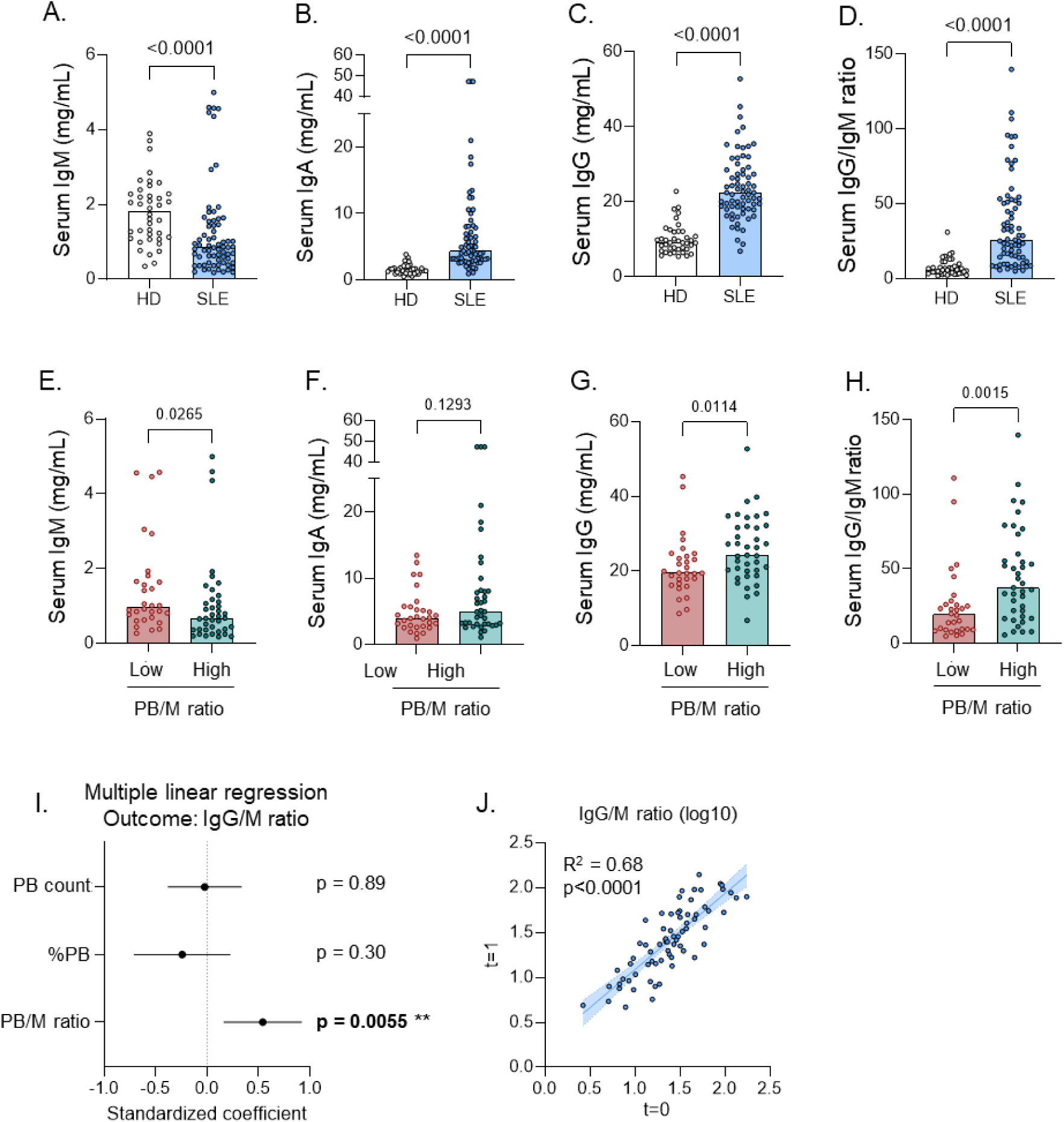
B cell hyperactivity results in hypergammaglobulinemia. Serum antibodies were measured in SLE patients (n=69) and healthy donors (n=40) using ELISA. SLE patients were split into two groups based on PB phenotype shown in Figure 1. A-H) Serum levels of total IgG, IgA, and IgM, and IgG:IgM ratio in SLE patients and healthy donors (A-D) and SLE patients split according to PB/CD27 group (E-H). I) Forest plot of multiple linear regression model with IgG:IgM ratio as outcome, and the Z-score of PB count, %PB, PB/M ratio, and cohort as predicting variables. J) Correlation of serum immunoglobulin levels in SLE patients measured at two timepoints ∼1 year apart (n=72). Each dot indicates an individual, and the bars represent the median (A-H,J). P values were obtained using Mann- Whitney test (A-H), multiple linear regression (I), or simple linear regression (J).

Importantly, serum immunoglobulin changes were less pronounced or not significant when patients were grouped based on the frequency of PB or the PB count (Figure S10A-H), or on the frequency of memory B cells (data not shown). Furthermore, in a multiple linear regression model with continuous values of the PB/M ratio, frequency of PB and PB count as predicting variables (thus independent from specific cut-offs used to group patients), only the PB/M ratio could significantly predict IgG:IgM ratios (Figure 5I). Like the PB/M ratio, levels of serum immunoglobulins significantly correlated over time, in particular for the IgG/IgM ratio (Figure 5J, Figure S10I-K).

Together, these data suggest that B cell hyperactivity in the IgG B cell compartment, reflected in a high PB/M ratio, results in stable hypergammaglobulinemia in SLE.

### B cell hyperactivity is specifically associated with the presence of Sm/RNP autoantibodies

Next, we wished to address the question whether specific ANA reactivities were increased in patients with a high PB/M ratio. In line with increased levels of total IgG, SLE patients with a high PB/M ratio had higher levels of total ANA-IgG (Figure 6A), and had a broader recognition profile in ANA (Figure 6B). Again, these differences were less pronounced or absent when grouping patients based on the frequency or absolute numbers of PB in circulation (Figure S11A-D). These findings were independently confirmed in a replication cohort (Figure S12A). Serum IgG reactivities to 8 nuclear antigens were measured by ELISA. Surprisingly, a strong increase in specific Sm/RNP reactivity was observed in patients with a high PB/M ratio (Table 1, Figure 6C-E). This association was only observed for Sm/RNP reactivity and not for the other ANAs. Again, these differences were much smaller or less significant when analyzing the PB frequency, Bmem frequency, or the absolute count of PB (Figure S11E-G, Table S5,S6). Multiple logistic regression for anti-Sm/RNP positivity with the different PB parameters revealed a significant relationship with the PB/M ratio. No such relationship was noted for the other PB parameters (Figure S11H).

**Figure 6:**
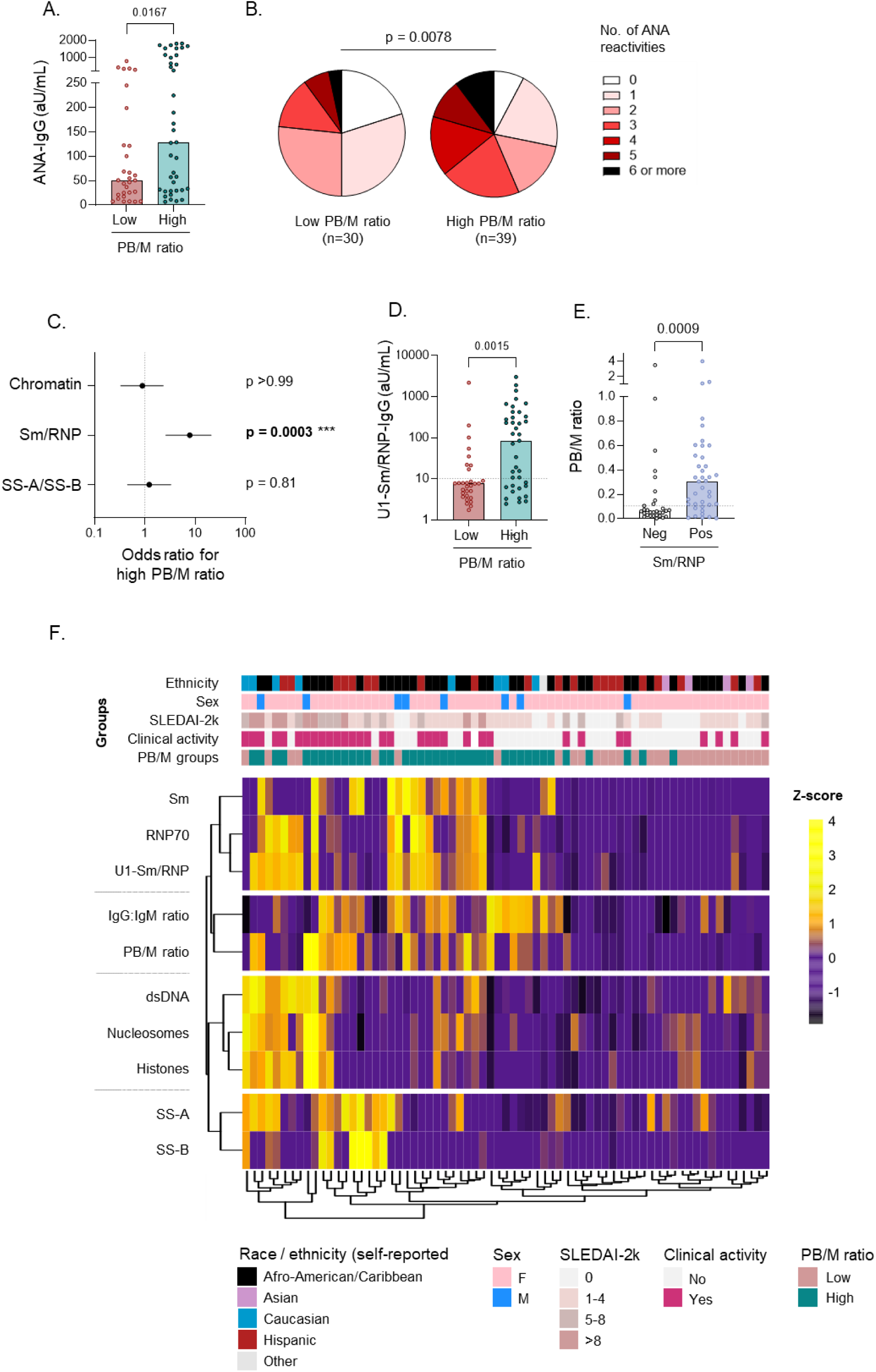
B cell hyperactivity is specifically associated with the presence of Sm/RNP autoantibodies. Serum autoantibodies were measured in SLE patients (n=69) using ELISA. Healthy donors (n=40) were used as negative controls. Autoantibody reactivities were categorized into three main antigen groups: chromatin (dsDNA, nucleosomes, histones); Sm/RNP (Sm, RNP70, U1-RNP complex), SS-A/B (SS-A, SS-B). SLE patients were split into two groups based on PB phenotype shown in Figure 1. A) Level of ANA-IgG in SLE patients split according to PB/M ratio. B) Number of IgG-ANA reactivities in SLE patient groups. C) Forest plots of odds ratios with 95% confidence intervals obtained with Fisher’s exact test for specific ANA reactivity with the indicated PB groups. Details are shown in Table 1. D) Level of U1-Sm/RNP-IgG in SLE patients split according to PB/M ratio. E) PB/M ratio in SLE patients split according to Sm/RNP-IgG reactivity. F) Heatmap with hierarchical clustering, of serological parameters and PB/M ratio in SLE patients on the rows, and individual patients in columns. Z-score was calculated for each row (parameter). Patient characteristics are shown in the bars above the heatmap. Each dot indicates an individual, and the bars represent the median (A,D,E). P values were obtained using Mann- Whitney test (A,B,D,E), or Fisher’s exact test (C).

**Table 1:**
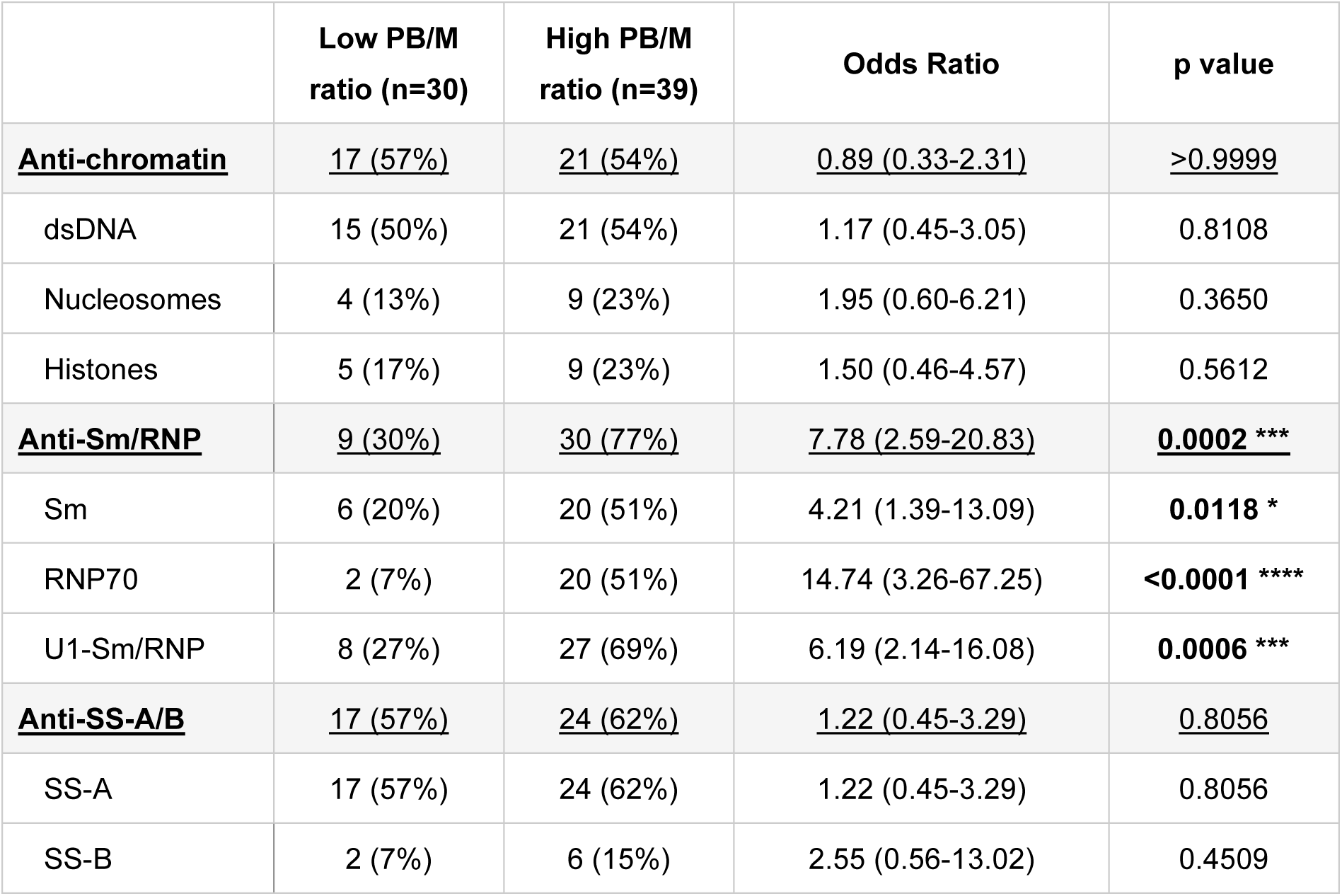
Frequency of ANA reactivities in SLE patients with low versus high PB/M ratios. Positivity of ANA-IgG for each indicated antigen was determined using ELISA. Anti-chromatin refers to all patients positive for 1 or more of the antigens dsDNA, Nucleosomes, and histones. Anti-Sm/RNP refers to all patients positive for 1 or more of the antigens Sm, and RNP70. Anti-SS-A/B refers to all patients positive for 1 or more of the antigens SS-A, and SS-B (all patients with SS-B were also positive for SS-A). P values were obtained using Fisher’s exact test.

|  | Low PB/M<br>ratio (n=30) | High PB/M<br>ratio (n=39) | Odds Ratio | p value |
| --- | --- | --- | --- | --- |
| **<u>Anti-chromatin</u>** | **<u>17 (57%)</u>** | **<u>21 (54%)</u>** | **<u>0.89 (0.33-2.31)</u>** | **<u>>0.9999</u>** |
| dsDNA | 15 (50%) | 21 (54%) | 1.17 (0.45-3.05) | 0.8108 |
| Nucleosomes | 4 (13%) | 9 (23%) | 1.95 (0.60-6.21) | 0.3650 |
| Histones | 5 (17%) | 9 (23%) | 1.50 (0.46-4.57) | 0.5612 |
| **<u>Anti-Sm/RNP</u>** | **<u>9 (30%)</u>** | **<u>30 (77%)</u>** | **<u>7.78 (2.59-20.83)</u>** | **<u>0.0002 \*\*\*</u>** |
| Sm | 6 (20%) | 20 (51%) | 4.21 (1.39-13.09) | **0.0118 \*** |
| RNP70 | 2 (7%) | 20 (51%) | 14.74 (3.26-67.25) | **<0.0001 \*\*\*\*** |
| U1-Sm/RNP | 8 (27%) | 27 (69%) | 6.19 (2.14-16.08) | **0.0006 \*\*\*** |
| **<u>Anti-SS-A/B</u>** | **<u>17 (57%)</u>** | **<u>24 (62%)</u>** | **<u>1.22 (0.45-3.29)</u>** | **<u>0.8056</u>** |
| SS-A | 17 (57%) | 24 (62%) | 1.22 (0.45-3.29) | 0.8056 |
| SS-B | 2 (7%) | 6 (15%) | 2.55 (0.56-13.02) | 0.4509 |

Both Sm reactivity and increased PB signatures have been previously associated with African-American (AA) race (11, 28, 29). Importantly, the association between the presence of a high PB/M ratio and Sm/RNP antibodies was statistically significant in non-AA SLE patients (OR: 10.31 (1.935 to 47.96); p=0.0047), displayed a trend for significance in AA SLE patients (OR: 4.56 (1,02-17,93; p=0.0623), and was replicated in an independent cohort (Table S7, Figure S12B-D). Therefore, a high PB/M ratio, that reflects B cell hyperactivity in SLE, is specifically associated with Sm/RNP IgG autoantibodies.

We summarized our findings in a heatmap in which Z-scores were calculated after log-transformation for serological parameters (all ANAs, and IgG:IgM ratio) and PB/M ratios in SLE patients (Figure 6F). Unsupervised hierarchical clustering analysis confirms the co-occurrence of ANA reactivities within antigen groups (Sm/RNP, chromatin, SS-A/B), which are known to frequently coincide within patients. Moreover, clustering of the parameters confirmed the correlation between PB/M and IgG:IgM ratio as measures of B cell hyperactivity. The PB/M ratio thus identifies a biologically relevant subgroup of SLE patients which is characterized by IFN-driven B cell hyperactivity and hypergammaglobulinemia, and displays frequent Sm/RNP reactivity and is more likely to have clinically active disease.

## DISCUSSION

In this study we identified a distinct phenotype in SLE patients which is characterized by a high PB frequency relative to CD27+ memory B cells. Patients with a high PB/M ratio also displayed other alterations within the B cell compartment. Whereas the lower frequency of CD27+ memory B cells was predominantly related to a decrease in unswitched memory B cells, CD27+ switched memory B cells and pre-PB were more activated in these patients, suggesting an increased propensity of switched memory cells to differentiate to PB. We found increased proliferation and a strong IFN signature in PB from these patients. In line with increased PB differentiation, class-switched antibodies in serum were increased suggesting the PB/M ratio signifies a hyperactive B cell response.

We observed that patients with a high PB/M ratio exhibit serological features of B cell hyperactivity, with a strong enhancement of class-switched immunoglobulins in serum and a broader ANA profile. Previous studies have reported hypergammaglobulinemia in SLE, in particular increased IgG, as well as low IgM levels (30–32). Our data confirmed these findings and revealed that low IgM and high IgG and IgA are particularly found in patients with a high PB/M ratio. Surprisingly, hypergammaglobulinemia and the IgG:IgM ratio only related to the PB/M ratio but not to the frequency of PB or PB count. One study reported a higher absolute PB count in patients with specific ANAs (dsDNA, Sm, SS-A and SS-B) in the absence of an effect on total IgG, IgA or IgM (14). Furthermore, changes in the PB signature in whole blood associated with changes in anti-dsDNA levels, but an association with hypergammaglobulinemia was not reported (33). Our results now link changes in the B cell subset composition with hypergammaglobulinemia in human SLE, providing evidence that B cell hyperactivity may directly lead to increased immunoglobulin production.

Patients with a high PB/M ratio displayed increased activation of switched B cells, in two distinct subsets; within the switched memory (CD27+IgD-) compartment, both the proportion of CD21^lo^ as well as their expression level of activation markers was increased. Within the DN (CD27-IgD-) B cell compartment, an increase in CD21^lo^ cells (DN2) was found as well, and these cells had increased expression of a more restricted set of activation markers. We found a strong correlation between the frequency of CD21^lo^ cells within switched memory B cells and DN cells. CD21^lo^ B cells have long been recognized as players in SLE and autoimmunity (21–23). Whereas several studies have shown that both DN2 and switched memory B cells have an altered phenotype in SLE, most focus has been placed on DN2 cells (23, 34). Several studies reported increased frequency of DN2 cells in SLE, in particular in patients with active disease (23, 35). In our study, an increased frequency of DN2 cells was less pronounced, and many patients with a high PB/M ratio or high %PB did not display an increase in DN2 cells. This suggests that PB expansion and hypergammaglobulinemia can be present without an expansion of DN2 cells. Different characteristics, such as different race/ethnicities, and clinical activity (relatively low disease activity and low frequency of lupus nephritis), between our cohort and previous cohorts on DN2 cells could explain the different findings on the frequency of DN2 cells .

In the context of infection and vaccination, resting memory B cells can give rise to both CD27+CD21^lo^ and CD27-CD21^lo^ B cells upon recall, even within the same clone (36). Both DN2 cells (CD27-CD21^lo^) and activated switched memory cells (CD27+CD21^lo^) have been reported to have a high propensity to differentiate into PB/PC (23, 37). Since we found increased activation within both subsets, hyperactivity within both CD27- as well as CD27+ B cells can be simultaneously present in SLE. Patients with a high PB/M ratio had increased frequencies of IgG1+ PB, and these cells displayed SHM rates comparable to those in switched (IgG1) memory B cells from SLE patients that we and others have observed (Figure 4H,I) (12, 38). Furthermore, CD45RB expression in PB was high, and similar to that of CD27+ memory B cells. This marker has been proposed to enable the tracking of activated B cells and PB derived from CD45RB+ memory B cells (27). Therefore, together these results point to activated switched memory B cells as a likely contributor to PB expansion in SLE. As such, the response has features of IFN-driven re-activation of memory B cells which have been previously generated through a GC response (39, 40). While IFN can enhance in vitro PB differentiation from memory B cells (41), IFN also plays an important role in GC B cell responses (42). The inability to study GC B cells in SLE patients is a limitation of our study.

Several studies have reported increased PB frequencies or counts in SLE (8, 11–14). Our data, that were replicated in an independent cohort, suggest that the ratio of PB to CD27+ memory B cells is more characteristic of SLE, than the PB frequency alone. Importantly, our results have shown that the PB/M ratio is associated with serological features of B cell hyperactivity.

Besides a correlation of the PB/M ratio with total immunoglobulins, we found a very strong association with Sm/RNP autoantibodies. Whether Sm/RNP antibodies arise as a consequence of enhanced PB differentiation from switched B cells or Sm/RNP positivity leads to a higher PB/M ratio remains to be determined. Sm/RNP antibodies can give rise to an IFN signature (43–45) which could subsequently cause increased PC differentiation from memory B cells (46). However, other studies report an association of IFN with other ANA reactivities or the breadth of the ANA response regardless of specificity (47, 48). Therefore, it remains to be determined whether Sm/RNP antibodies give rise to B cell hyperactivity and a high PB/M ratio, or the other way around. It is also plausible that they are part of a feedforward loop in which they amplify each other.

Higher IgG levels (49), a higher PB signature (11), Sm/RNP antibodies (28, 29), and higher disease activity (50, 51) have been associated with African-American ancestry in SLE patients, though these features can be present in SLE patients of any descent. In our study, the association between PB/M ratio and Sm/RNP reactivity was replicated in an independent European cohort, and was present in both AA and non-AA patients from the US cohort. Thus, though the hyperactive B cell phenotype, characterized by a high PB/M ratio, is more commonly observed in AA patients, it is also present in non-AA patients, and as such, represents a distinct biological patient phenotype regardless of race. The relationship of a high PB/M ratio with disease activity suggests that B cell hyperactivity may play a role in disease pathogenesis and may thus guide the tailoring of B cell targeted therapies. The strong association with Sm/RNP reactivities indicates that serum autoantibody profile in combination with determining the B cell distribution (PB and CD27+ memory B cells) can be used to determine the degree of B cell hyperactivity and may be used for patient stratification in clinical trials, in particular those global B cell activation, PB differentiation and type I IFN.

## Acknowledgements

This work was supported by the Bontius foundation (JS), LUMC funding (internal MSCA Seal-of-Excellence 2020 regulation; JS), an Investigator-Initiated Research award ASPIRE provided by Pfizer (JS), a fellowship from the Arthritis National Research Foundation (YA-F) and NIH grants P01AI148102 and U19AI144306.

## Author contributions

H.J.v.D. contributed to the design of the study, acquisition, analysis and interpretation of data and the draft of the manuscript. Y.A.F. contributed to the design of the study, the acquisition, analysis and interpretation of data and the revision of the manuscript. A.L.D contributed to the design, acquisition and analysis of data. T.W.J.H., M.M., and C.A. contributed to the acquisition and interpretation of data and the revision of the manuscript. R.E.M.T and B.D. contributed to the design of the study, the interpretation of data and the revision of the manuscript. J.S. contributed to the design of the study, the acquisition, analysis and interpretation of data, and the draft of the manuscript.

## METHODS

### SLE patients and healthy subjects

<u>Cohort 1:</u> Heparinized blood from 72 SLE patients (of which 36 were previously reported on) (20) and 14 healthy subjects was collected from The Feinstein Institute for Medical Research Rheumatology Specimen and Clinical Data Bank (Manhasset, NY, USA), and PBMCs were obtained using standard Ficoll procedure. Serum was obtained from SLE patients at the time of PBMC isolation and was available for 69 patients. From 24 patients, a second sample of heparinized blood, was obtained to determine the stability of B cell phenotypes. Timepoints were more than 1 month apart (median 10; range: 1.5-26). SLE diagnosis was made clinical and fulfilled the 1997 revised ACR classification criteria (3). SLE patients were excluded from the study if they received belimumab, rituximab, or cyclophosphamide 12 months prior to the study. Clinical data, including SLEDAI-2k and c-SLEDAI scores (52), were collected from the same timepoint as B cell analysis. Clinically active patients were defined as patients with a c-SLEDAI above zero.

<u>Cohort 2</u>: A replication cohort consisted of 77 SLE patients from the Rheumatology outpatient clinic at the Leiden University Medical Center (Leiden, The Netherlands) from which serum was available. Of these, B cell phenotype obtained in freshly isolated PBMCs was measured in 37 SLE patients and in frozen PBMCs in 15 patients, with 5 patients measured in both fresh and frozen PBMCs obtained at the same time. Complete disease activity data was available for 17 of these SLE patients. B cell phenotype was also measured in fresh PBMCs from 21 healthy subjects, and in frozen PBMCs from 10/9/8? Healthy subjects. One patient was excluded for IgA analysis as it was deemed IgA-deficient. Serum from 40 healthy donors were used to calculate the normal values and cut-offs for ELISA, and were obtained from the LUMC Voluntary Donor Service.

Patient characteristics from both cohorts are described in Table S1 and S2. The study with SLE patients and healthy subjects from the USA was approved by the Northwell Health Institutional Review Board, and the study with PBMCs from SLE patients and healthy subjects from Leiden were approved by the Leiden-Den Haag-Delft Medical Ethical Committee. The use of serum samples from healthy subjects was approved by the LUMC-biobank organization. All subjects gave written informed consent.

### Flow cytometry

PBMCs were stained for surface markers using fluorescent antibodies diluted in HBSS + 2% FCS (Cohort 1) or PBS containing 1% BSA (Cohort2). eFluor 506–labeled fixable viability dye (eBioscience) was added during staining with cell surface antibodies. For spectral flow cytometry, antibody cocktails were prepared using Brilliant Stain Buffer (BSB) Plus (BD) and monocyte blocker (Biolegend) was added to prevent aspecific binding of tandem dyes. For samples with surface staining alone, cells were washed and left unfixed or were fixed with 1% PFA. For samples with intracellular staining, cells were washed and fixed and permeabilized with Foxp3 transcription factor fixation/permeabilization kit (eBioscience) according to manufacturer’s instructions. Antibodies used in this study are shown in Supplementary table S8 and S9.

Conventional flow cytometric acquisition was performed on a Fortessa (BD). FACS sorting was performed on FACS Aria or Aria Sorp sorters (BD). Analysis of conventional flow cytometry and FACS sorting was performed using FACS Diva (BD) and FlowJo software. Spectral flow cytometric acquisition was performed on a 5-laser Aurora (Cytek).

### Spectral flow cytometry data analysis

Raw spectral data was unmixed using Spectroflo software (Cytek), after which unmixed fcs files were analysed in OMIQ. A batch normalization control pooled PBMC sample was taken along in each experiment (4 experiments with fresh PBMCs, 1 experiment with frozen PBMCs). Live B cells were gated, after which expression of CD19, CD20, CD21, CD24, CD27, CD38, IgD, IgM, IgG and IgA across each experiment was normalized using CytoNorm (53). Subsequently, live B cells were clustered with FlowSOM (elbow metaclustering) using CD19, CD20, CD21, CD24, CD27, CD38, IgD, and IgM as input features. Expression of markers was analysed in UMAP and heatmaps to identify known B cell subsets (40). Fifteen FlowSOM metaclusters were obtained, from which 2 clusters were very similar in expresson patterns and were together designated as resting naive. One small cluster (<1%) was identified as non-B cells and excluded from downstream analysis, leaving 13 final B cell clusters. Using the same features as in FlowSOM, 500k subsamples cells across all samples were projected onto UMAP for visualization (neighbors = 100, mindist = 0.4). Cell counts and median fluorescence intensities (MFI) for each cluster and each sample were exported from OMIQ for further processing and statistical analysis. BST2 expression was normalized by subtracting the MFI of control cells (stained without BST2) and divided by the BST2 MFI of all B cells in the batch control PBMC sample.

Clustered heatmaps were made using scaled data obtained from OMIQ. Heatmaps for cluster identification was performed using median expression levels within subsets, Euclidean distance and Ward linkage in OMIQ. Hierarchical clustering displayed on heatmaps and PCA analysis for activation marker expression within subsets and patients were performed in R using pheatmap and prcomp respectively. Trajectory analysis was performed using Wanderlust in OMIQ using k=15 and 250 waypoints. After defining a small population of the most immature Tr1 cells, Wanderlust placed cells on a one-dimensional developmental trajectory, culminating in the most terminally differentiated PB cluster.

### scRNAseq library preparation and sequencing

PB (CD3/14/FVD-CD19+CD27++CD38++) were isolated by FACS sorting, washed in HBSS or PBS + 0.04% non-acetylated molecular grade BSA (Thermo), centrifuged at 1000xg for 5 min, and resuspended in the same buffer. Cell suspensions were processed through the Chromium Single-Cell 5′ RNA-seq system (10x Genomics), according to the manufacturer’s protocols. In the first experiment, cells from 4 patients in Cohort 1 were run on separate lanes using Chromium Single Cell 5’ Library & Gel Bead Kit v1.1. In a second experiment, cell hashtags (Biolegend Totalseq antibodies) were used to multiplex frozen cells from 5 patients from Cohort 2 on a single lane using Chromium Single Cell 5’ Library & Gel Bead Kit v1.1 as well as the Single Cell 5’ Feature Barcode Library Kit (10x Genomics). After cDNA amplification, BCR target enrichment was performed using the Chromium Single Cell V(D)J Enrichment Kit for Human B cells (10x Genomics). Sequencing of libraries from the first experiment was performed on a HiSeq system (Illumina) with 2×150bp reads. Sequencing of the second experiment was performed on a partial lane of the illumina NovaSeq6000 at a 151-16-9-151 bp run configuration. The target sequence depth was 25K reads/cell for Gene expression, 5K reads/cell for BCR, and 2.5K reads/cell for Hashtag oligo. Sequencing saturation was > 70% in each sample.

### scRNAseq data analysis

Raw sequencing data was demultiplexed and aligned to the human genome (GRCh38-2020-A) using Cellranger software v4.0.0 (Experiment 1) and v5.0.1 (Experiment 2) (Chromium 10x). Analysis was performed with the Seurat v5.0.1 R package(54). Ig and TCR variable genes were removed to avoid bias in clustering/gene expression analysis by unique V gene usage. BCR repertoire was analysed separately (see below). Contaminating cells (non-PB) were removed following celltype prediction through the SingleR package using the Monaco immune reference dataset (13/5.5% removed) (55, 56). Low quality cells were removed based on a low number of detected genes and a high frequency of mitochondrial reads per cell (Experiment 1: >200 genes and <10% mt; Experiment 2: >2000 genes and <8% mt). Samples from run 2 were demultiplexed with the HTOdemux function in Seurat, followed by removal of Negative/ambiguous and Doublet cells (4.3%) (57). In experiment 1, a small fraction of cells with the same barcode (<2%) was present in samples from different lanes (likely due to contamination), these were removed by retaining the cell with the highest read count. Final cell numbers for analysis were n=1290 cells from Experiment 1 and n=1197 cells from Experiment 2.

After pre-processing each Experiment individually, experiments were combined in Seurat, and integrated analysis was performed using anchor-based canonical correlation analysis (CCA) in the function IntegrateLayers. UMAP and FindNeighbors was performed using the first 20 integrated CCA dimensions. Clusters were identified by shared nearest neighbour (SNN) modularity optimization using FindNeighbors with CCA as reduction and 1:20 dimensions as well as FindClusters with resolutions ranging from 0.1 to 0.5 in 0.1 increments using FindClusters. By visual inspection and analysis of differentially expressed genes, and consistency across both experiments, clustering with a resolution of 0.1 (resulting in 2 clusters) was determined to best reflect distinct PB subsets. UMAP was generated using RunUMAP (n.neighbors = 100, min.dist = 0.4, spread = 1) and revealed good integration of both experiments.

Data were combined with JoinLayers for differential gene expression analysis. Data were normalized by the NormalizeData() function and scaled by the ScaleData() function. Covariate neighborhood analysis (CNA) was performed within Seurat using the rcna package (58) and visualized in UMAP space to analyse which neighborhoods were most associated with patient groups. Differential gene expression analysis was performed using FindMarkers (test.use = MAST, latent.vars = Experiment, logfc.threshold = 0.25). FindMarkers was performed twice, comparing the two identified PB clusters and comparing the two patient groups. One male patient in the Low PB/M group was excluded for comparison of the two patient groups as it led to identification of many male-specific transcripts in this group. This patient was included for subsequent analyses. Significant marker genes (adjusted p value<0.05) were exported in csv files and pathway analysis was performed using DAVID for gene ontology (GO) terms (biological pathways; BP). The resulting functional annotation chart was used in Cytoscape Enrichment Map for visualization of pathway enrichment (using cutoffs: FDR Q-value: 0.05; Overlap: 0.5; Test used: Overlap index) (59). Gene set enrichment analysis (GSEA) was performed with the ranked list of significant marker genes between groups using the GSEA function of the package ClusterProfiler. Source databases for GSEA were obtained from the GSEA molecular signature database (MsigDb) (60), and included hallmark cytokine gene sets, IFN-regulated transcription factors from dataset C. An additional dataset obtained from PBMCs stimulated with various IFNs and cytokines was used to generate a more detailed IFN signature (61)

### 5’ RACE PCR

ANA+ B cells were stained as described (8), and ANA+ naïve (CD19+CD27-IgD+), double-negative (CD19+CD27-IgD-), and memory B cells (CD19+CD27+) were sorted from PBMCs of SLE patients used in scRNAseq experiment 2 using a FACS Aria (BD). 0.4-35k cells per population were lysed in RLT buffer (Qiagen) with 10 uL/mL 2-ME (Merck) after which RNA was isolated using RNeasy Microprep. cDNA was generated and amplified by incubating 8 uL RNA with Oligo-dT30VN (2.5 uM) and dNTPs (2.5 uM) for 3 min. at 72°C. All primers were ordered from IDT and sequences are provided in Table S10. Subsequently, first-strand buffer (Takara), Betaine BioUltra (1 M, Sigma-Aldrich), DTT (5 mM, Takara), recombinant RNase inhibitor (1 U/uL, Takara), SMARTScribe reverse transcriptase (5 U/uL, Takara), Template-Switching Oligo (TSO, 1 uM) was added per sample. Samples were incubated for 90 min at 42°C, followed by 10 cycles of 2 min at 50°C and 2 min at 42°C, followed by incubation for 15 min at 72°C.

5’ RACE PCR products for Ig heacy chains (IgM, IgG, IgA) were generated using an adapted protocol for low cDNA amount based on the Anchoring Reverse Transcription of Immunoglobulin Sequences and Amplification by Nested PCR (62, 63). 2 μL of cDNA was added to a combination of Phusion Flash High-Fidelity PCR Master Mix, nuclease-free H_2_O, SA forward primer (200 nM, SA.PCR_2) and either one of Ig-specific reverse primers (40 nM, IgM.PCR, IgG.PCR, IgA.PCR). Mixtures were incubated for 2 min at 98°C, followed by 40 cycles of 1 sec at 98°C, 15 sec at 69°C and 15 sec at 72°C, and final extension for 1 min at 72°C. PCR products were purified using Qiaquick PCR purification kit (Qiagen) according to manufacturer’s instructions. Subsequently, 4 uL of each sample was barcoded by adding a mix of Phusion Flash High-Fidelity PCR Master Mix, nuclease-free H_2_O, one of the SA forward barcode family primers (200 nM) and one of the IgH-specific reverse barcode family primers (200 nM). Mixtures were incubated for 2 min at 98°C, followed by 10 cycles of 5 sec at 98°C, 15 sec at 65°C and 30 sec. at 72°C, and final extension for 5 min at 72°C. Samples were loaded on a 1% Agarose gel, followed by excision of the bands and purification using the Nucleospin Gel & PCR cleanup kit (Bioké). Different isotypes from the same sample were pooled. PCR products were quantified using the Qubit dsDNA Quantitation kit high sensitivity (Thermo Fisher), and samples were pooled in relative amount to their original cell number. Pooled PCR products were sequenced on a 8M SMRT cell (PacBio Sequel II) by the Leiden Genome Technology Center.

### BCR repertoire analysis

BCR repertoire was analysed using the scRNAseq data and the 5’RACE data. For the first scRNAseq dataset, filtered contig sequences for V(D)J were generated from the Gene Expression data using Trust4 (64). For the second scRNAseq dataset, enriched V(D)J libraries were aligned using Cellranger software to generate filtered contig sequences. For the 5’ RACE PCR, raw sequencing reads were processed using pRESTO in Python. Sequences were trimmed using trimqual (Quality score threshold 50 and window size 5), after which sequences with a minimum length of 400 were selected. High quality sequences (quality score ≥90) were selected. Sample barcodes were assigned using MaskPrimers. Duplicate sequences were concatenated using CollapseSeq. C regions were assigned using MaskPrimers. Sequences with a minimum read count of 2 were retained using SplitSeq.

The BCR sequences from all three datasets were used as input for IMGT High V-quest (version 1.9.4; IMGT/V-QUEST reference directory release: 202405-2 F+ORF+ in-frame P - With all alleles; with search for insertions and deletions; other settings default).

Output from IMGT High V-quest was subsequently processed using the Immcantation pipeline. First, AIRR databases were created from the IMGT input and output files using the function MakeDb.py imgt in pRESTO (Python). Clonal analysis was done using SCOPER in R. Productive sequences were selected (>99%). Nucleotide Hamming distance was calculated and normalized by junction length using distToNearest. Hamming distance threshold was calculated using the density method in findThreshold. The resulting threshold of 0.145 was used to identify clones using hierarchicalClones. Clonal abundance and diversity was determined using Alakazam after downsampling to 100 cells per patient (3 patients excluded). Results were confirmed using several degrees of downsampling. Clonal diversity was calculated using alphaDiversity at q=0 to q=8. V gene usage was calculated using the Alakazam package. Germline sequences were created using the createGermlines function in Dowser after which somatic hypermutation was calculated using the observedMutations function in the SHazaM package.

### RNA isolation and qPCR

CD19+CD20loCD27hiCD38hi PBs were sorted from PBMCs of SLE patients using a FACS Aria (BD). 1- 30k cells per population were lysed in RLT buffer (Qiagen) with 10 uL/mL 2-ME (Merck) after which RNA was isolated using RNeasy Microprep. cDNA was synthesized using iScript (Biorad). qPCR was performed after pre-amplification using TaqMan PreAmp Master Mix (Thermo) and the Taqman assays mentioned below. qPCR was performed with TaqMan Fast Advanced Master Mix (Thermo) and multiplexed VIC and FAM Taqman assays (all from Applied Biosystems/Thermo): POLR2A VIC-MGB (Hs00172187_m1), ACTB VIC-MGB (Hs01060665_g1), IFI6 FAM-MGB (Hs00242571_m1), BST2 FAM-MGB (Hs00171632_m1). qPCR was run on a CFX Opus machine (Biorad) using recommended cycling conditions. Relative expression in each sample was determined as the 2^(delta Ct) using the average of both housekeeping genes (POLR2A and ACTB).

### ELISA

384-well Clear Flat Bottom Polystyrene High Bind Microplates (Corning) were coated overnight at 4C with 1 µg/ml UltraPure™ Salmon Sperm DNA Solution (Thermo), 0,5 µg/ml non-recombinant bovine nucleosomes, 0,5 µg/ml non-recombinant bovine histones, 0,8 µg/ml non-recombinant bovine Sm, 0,6 µg/ml recombinant human U1snRNP-68/70kDa, 0,8 µg/ml non-recombinant bovine Sm/RNP70 complex, 0,5 µg/ml 60-kDa recombinant human SS-A or 1,1 µg/ml recombinant human SS-B (all from Diarect) diluted in PBS.

All washing steps were performed by washing 4 times with 0.05% Tween20 (Sigma) in PBS using the Berthold Zoom HT LB 920 Plate Washer. Plates were washed and subsequently blocked with 1% BSA (Sigma) and 50mM Tris (Roche, pH = 8,0) in PBS for 1 hour at RT. Subsequently, the plates were washed and the wells were incubated with serum, diluted 1:200 in PBS/0.05% Tween20/1% BSA/Tris (pH = 8,0), for 1 hour at RT. After the incubation, the ELISA plates were washed and incubated with 1 μg/ml anti-IgG horse-radish peroxidase (Bethyl) and, subsequently, washed and incubated with ABTS (Sigma-Aldrich) with 1:2000 hydrogen peroxide (Merck), All plates were measured at 415nm on the SpectraMax i3x Multi-Mode Microplate Reader. ANA standards consisted of pooled sera from positive SLE patients for each reactivity. In-house ANA ELISAs were validated using SLE serum samples with known reactivities obtained from the anti-ENA SLE profile 2 ELISA kit (Euroimmune) or clinical diagnostic assays (Figure S13). All ELISAs reached a positive and negative predictive value of >0.8.

Total IgG, IgM and IgA ELISAs were performed by coating 384-well Clear Flat Bottom Polystyrene High Bind Microplates with 10 µg/ml goat anti human IgG/IgM and IgA Fc (Bethyl) diluted in mQ water with 0,1 M Na_2_CO_3_/ NaHCO_3_ (Merck, pH = 9,6) for 1 hour at RT. Same protocols were used as for ANA ELISAs except serum samples were diluted 800.000x for total IgG, 80.000x for total IgA, and 20.000x for total IgM. Commercial standards with quantified IgM and IgA levels (Bethyl) or IgG (Southern Biotech) were used for quantification.

ANA and total Ig levels were expressed as aU/ml or mg/mL, respectively, based on the intrapolation from the standards using a 5-parameter logistic curves. For autoantibodies, samples were considered positive when they were above the cut-off, determined by the mean value + 4x std of 40 healthy donors. One and two healthy donors were excluded for determination of the cut-off for anti-SS-A and anti-Sm, respectively, because they were above the cut-off of the rest of the healthy donors, and had a large impact on the sensitivity of the assay determined by commercial ELISA.

### Statistics

Statistical analysis was performed in GraphPad version 10.2.3. P-values were considered statistically significant if they were below 0.05. Fisher’s exact test was used to analyse categorical data. For continuous data, Mann Whitney U tests were used to compare two groups, except qPCR which was analysed using Student’s t-test after log-normalization. Kruskal-Wallis was used to compare multiple groups, with false discovery rate (FDR) as posthoc test calculated using the Two-stage linear step-up procedure of Benjamini, Krieger and Yekutieli. Two-way ANOVA was used for statistical analysis of grouped data, with FDR as posthoc test calculated using the Two-stage linear step-up procedure of Benjamini, Krieger and Yekutieli. Correlations were analysed using simple linear regression. For multiple linear and logistic regression, continuous variables were log-transformed and converted to Z-scores for standardization.

Cut-offs of %PB and PB/M ratio were based on data from 14 healthy donors from Cohort 1, 21 donors from Cohort 2 (fresh) and 8 donors from Cohort 2 (frozen), calculated using Q3 + 1.5*IQR. One healthy donor from the Feinstein Institute and 2 from the Leiden cohort were excluded for calculation of the cutoffs as it exceeded the cut-off and were thus deemed outliers. These subjects were only excluded for determination of the cut-offs but remained present for statistical comparison of healthy subjects versus SLE patients. The resulting cutoffs were 2.75 for %PB, 0.105 for PB/M ratio on fresh cells, and a PB/M ratio of 0.035 for frozen cells. The latter was 3-fold lower on frozen cells compared to fresh cells due to specific loss of PB after freezing (Figure S4A). Using 5 SLE samples with paired fresh and frozen data, the PB/M ratio was also decreased approximately 3-fold (Figure S4B). For absolute PB count, healthy donor count data was unavailable, therefore these groups were divided based on the median of PB counts in SLE patients in the Feinstein cohort (1.91 cells/uL).

## SUPPLEMENTARY DATA

**Figure S1:**
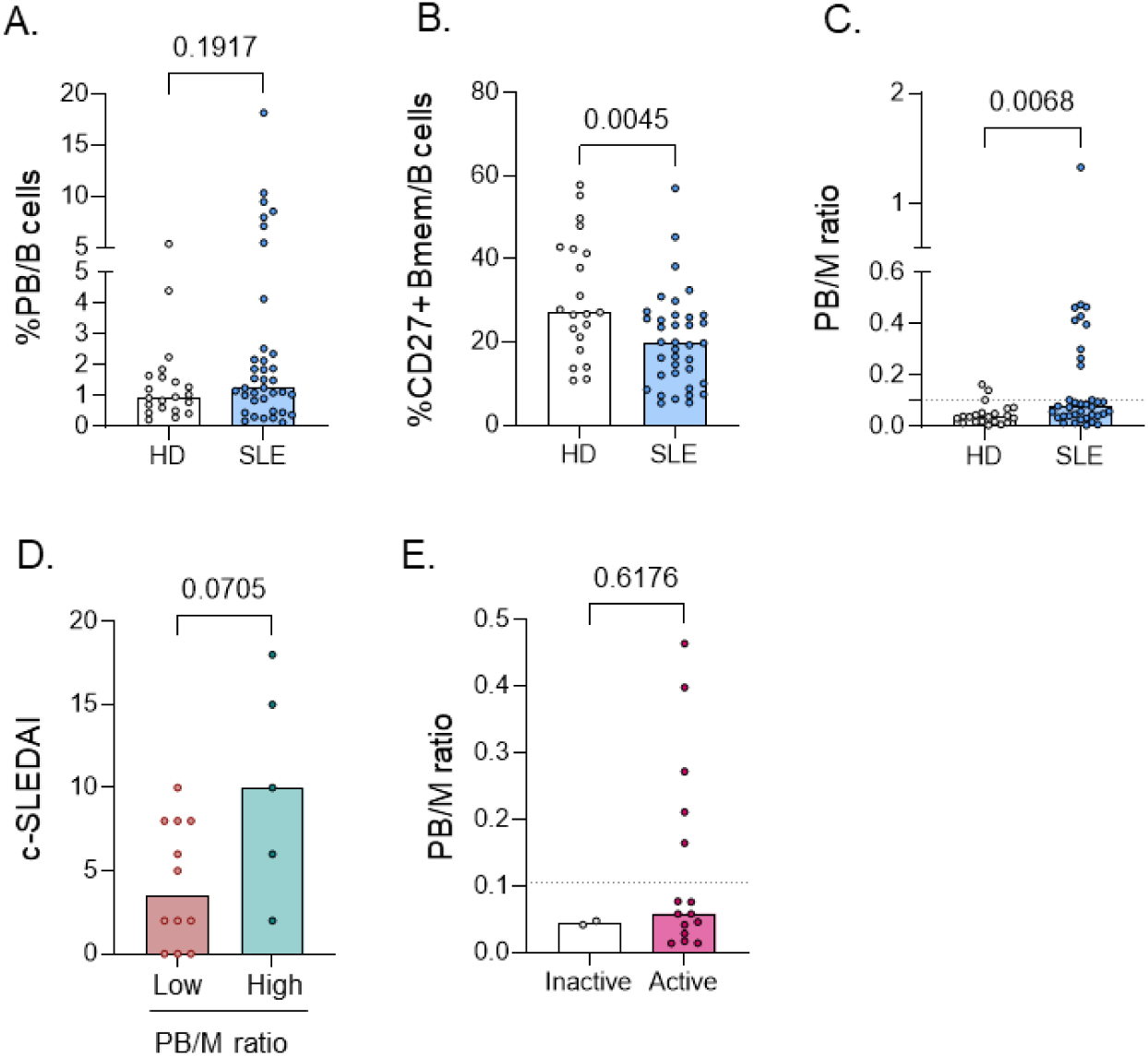
A high PB to memory B cell ratio characterizes a subgroup in SLE in cohort 2. B cell phenotypes of SLE patients (n=37) and healthy donors (n=21) were analysed using flow cytometry. Clinical data was available for 17 of these SLE patients. A-C) %PB, %CD27+ Bmem among total B cells, and PB/M ratio. D) c-SLEDAI in SLE patients with a low versus high PB/M ratio. E) PB/M ratio in patients with clinically inactive versus active (c-SLEDAI>0) disease. Each dot indicates an individual, and the bars represent the median. P values were calculated using Mann- Whitney test.

**Figure S2:**
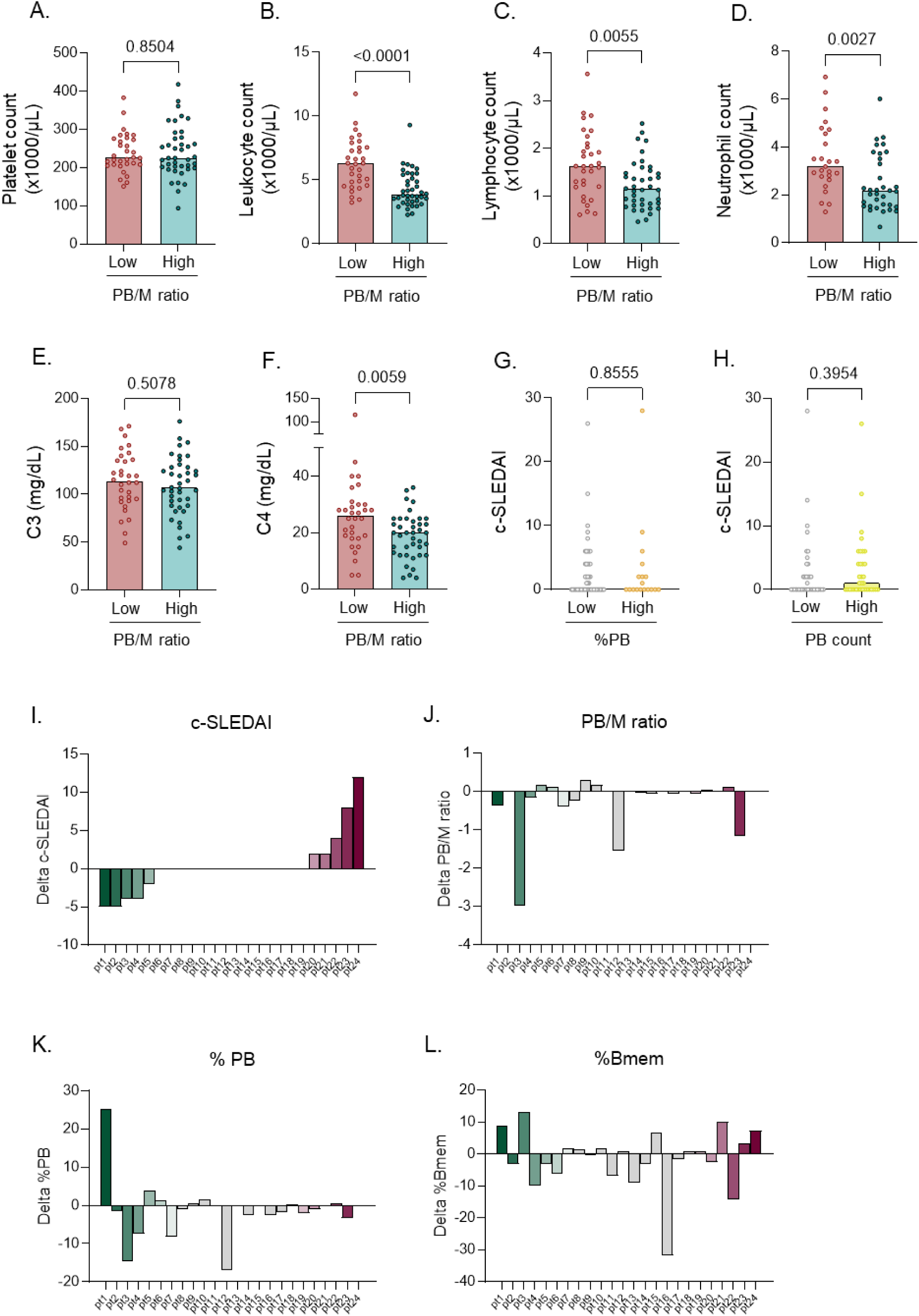
Additional analysis of disease activity in SLE patient groups. B cell phenotypes of SLE patients (n=72) and healthy donors (n=14) were analysed using flow cytometry. A-F) Differential blood cell counts and complement levels in SLE patients with a low versus high PB/M ratio. G,H) c- SLEDAI in SLE patients with a low versus high %PB or PB count. I-L) c-SLEDAI change over time and lack of association with changes in B cell subset percentages in SLE patients whose B cell phenotype was measured twice (n=24). Patients were sorted based on delta c-SLEDAI (I), after which the delta PB/M ratio (J), delta %PB (K), and delta %CD27+ Bmem were plotted in the same order. Each dot indicates an individual, and the bars represent the median. P values were calculated using Mann- Whitney test (A-H).

**Figure S3:**
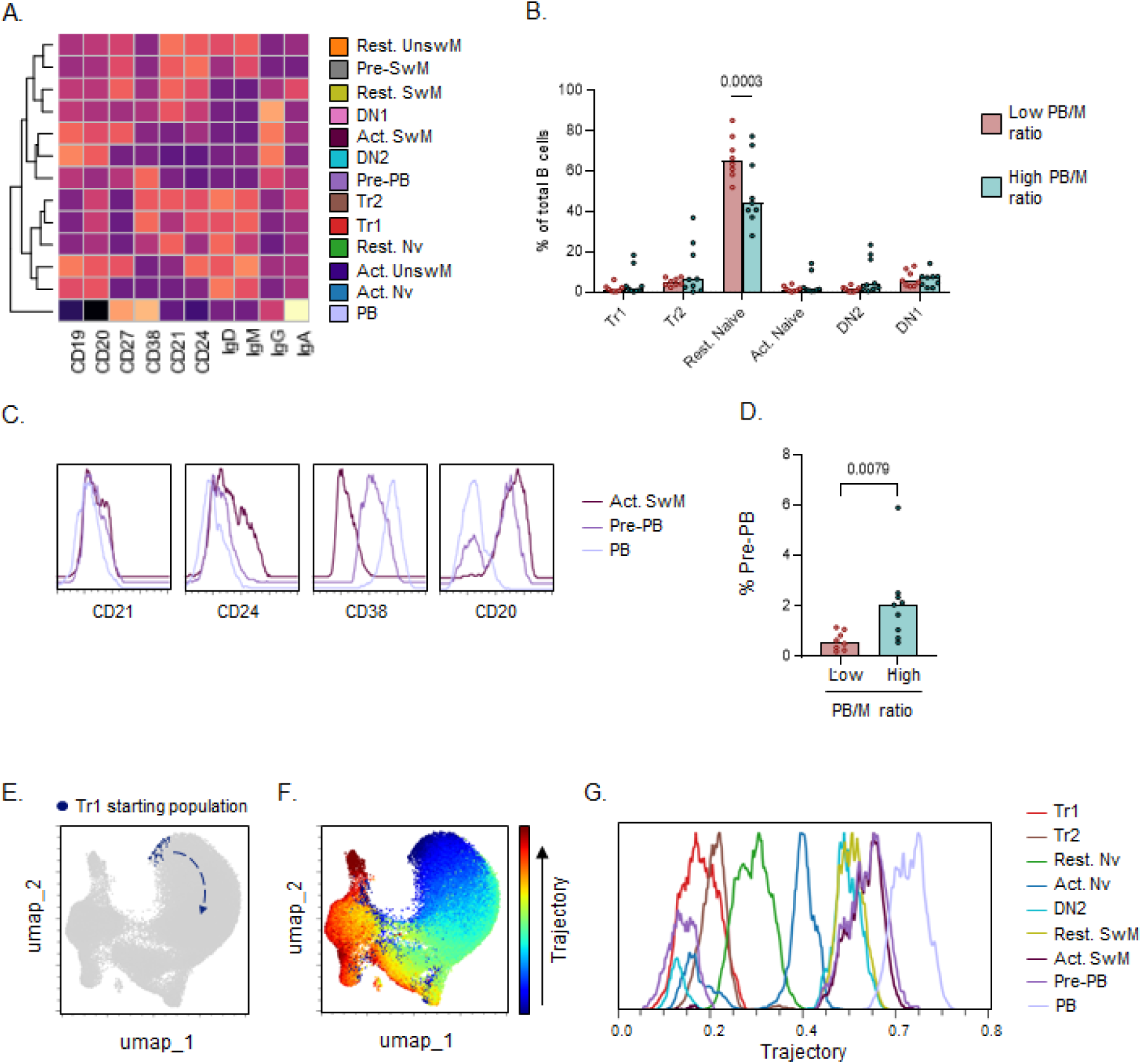
Additional spectral flow cytometry phenotyping in fresh PBMCs. High dimensional spectral flow cytometry was used for detailed B cell phenotyping within fresh PBMCs from SLE patients (SLE; n=17). A) Heatmap showing expression levels of markers (columns) for each FlowSOM cluster (rows). Z-score was calculated using the median expression among all samples. B) % of CD27- B cell populations SLE patients with a low versus high PB/M ratio. C) Histograms of key markers defining the Pre-PB cluster. D) % Pre-PB SLE patients with a low versus high PB/M ratio. E) Trajectory analysis using Wanderlust. Cells indicated as Tr1 in left UMAP were the starting population. F) Resulting trajectory projected on UMAP. G) Location of main B cell subsets on the Wanderlust trajectory, displayed as a histogram of maximized cell counts per cluster. Each dot indicates an individual, and the bars represent the median. P values were calculated using Mann- Whitney test (D), or Two-way ANOVA with FDR posthoc test (B).

**Figure S4:**
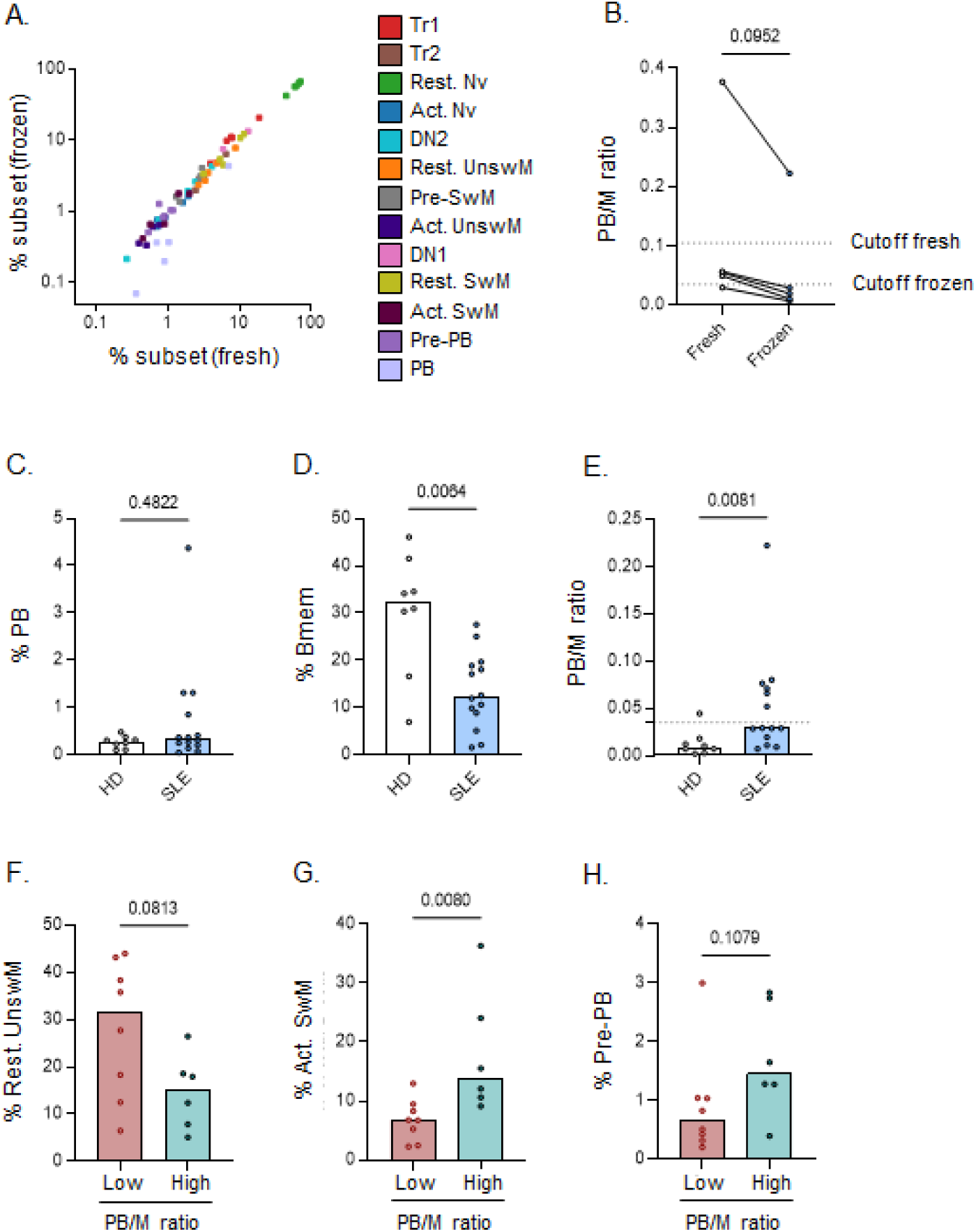
Replication of spectral flow cytometry phenotyping in frozen PBMCs. High dimensional spectral flow cytometry was used for detailed B cell phenotyping within frozen PBMCs from SLE patients (n=15) and healthy donors (HD; n=10). For 5 SLE patients, paired analysis of fresh PBMCs was also performed. A) Correlation of the percentage of each cluster in paired frozen and fresh samples, showing only the PB cluster is affected by freezing. B) Paired analysis of the PB/M ratio in fresh and frozen samples. The cutoff for each dataset is indicated and was based on the distribution in healthy donors (Q3+1.5*IQR). C-E) %PB, %CD27+ Bmem, and PB/M ratio in healthy donors compared to SLE patients. F-H) %Rest UnswM, Act SwM, Pre- PB SLE patients with a low versus high PB/M ratio. Each dot indicates an individual, and the bars represent the median. P values were calculated using Mann- Whitney test (C-H).

**Figure S5:**
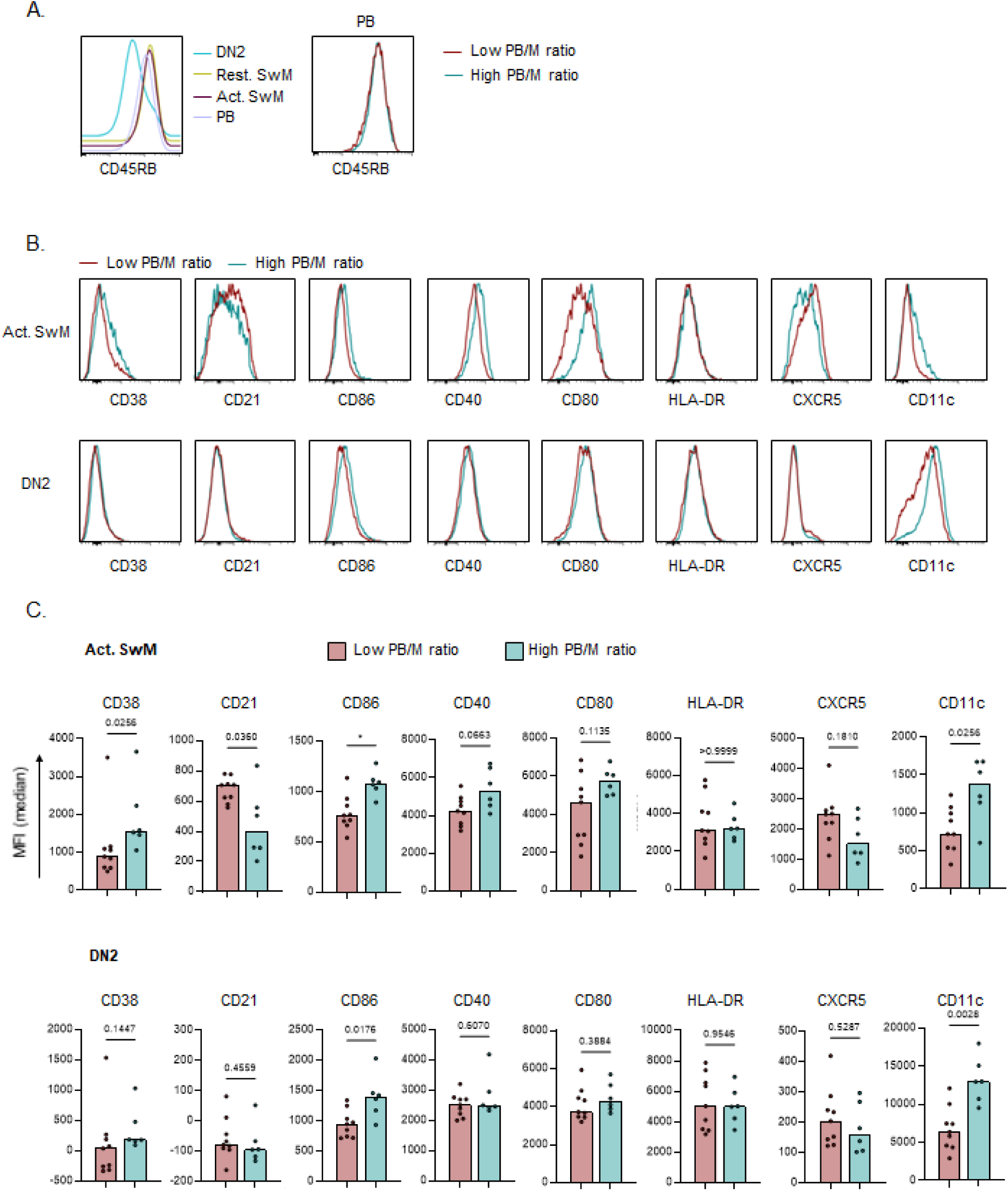
Additional spectral flow cytometry phenotyping in frozen PBMCs. High dimensional spectral flow cytometry was used for detailed B cell phenotyping within frozen PBMCs from SLE patients (n=15) and healthy donors (HD; n=10). A) CD45RB expression in several switched B cell subsets (left) and comparison of CD45RB expression in PB from SLE patient groups. B) Histograms showing expression of activation markers within Act SwM (top row) and DN2 cells (bottom row) in SLE patients with a low versus high PB/M ratio. C) Expression level of activation markers between the two patient groups in Act SwM (top row) and DN2 cells (bottom row). Each dot indicates an individual, and the bars represent the median. P values were calculated using Mann- Whitney test (C).

**Figure S6:**
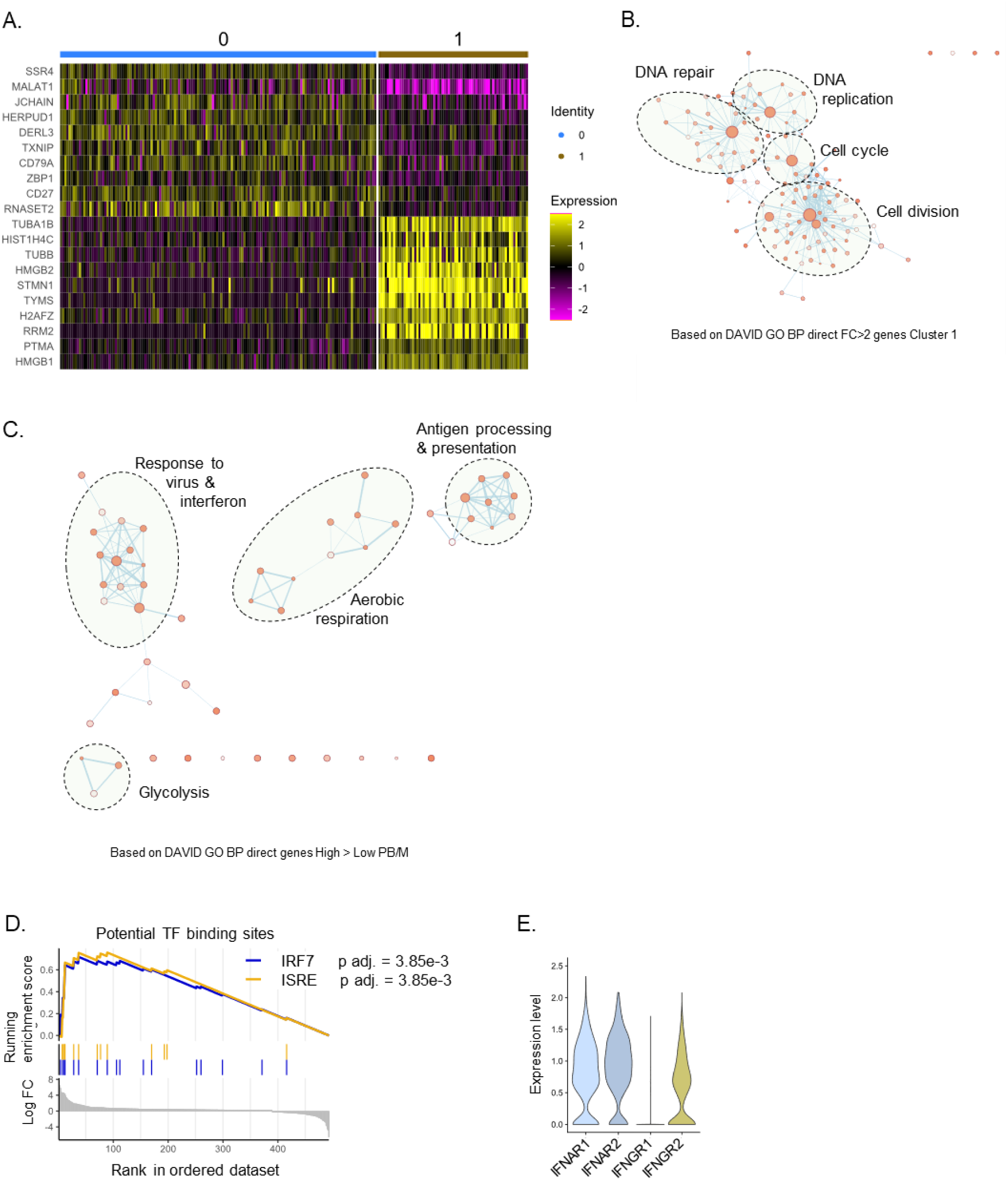
Increased proliferation and IFN signature underly PB expansion in patients with a high PB/M ratio. scRNA-seq was performed on sorted PB from 9 SLE patients, in 2 independent experiments. A) Heatmap of top 10 differentially expressed genes (adjusted p value <1*10e-10, highest fold change) between patients with a low versus high PB/M ratio. 150 cells from cluster 0 and 75 cells from cluster 1 are shown. B) Clustering of enriched pathways in cluster 1 determined using DAVID functional enrichment analysis and visualized using enrichmentMAP. C) Clustering of enriched pathways in patients with a high PB/M ratio as in B. D) GSEA for regulatory target gene sets and predicted transcription factor binding sites of IFN-regulated factors using differentially expressed genes (adjusted p value <0.05; ordered by log fold change) between the two patient groups. E) Expression level of IFN receptor subunits.

**Figure S7:**
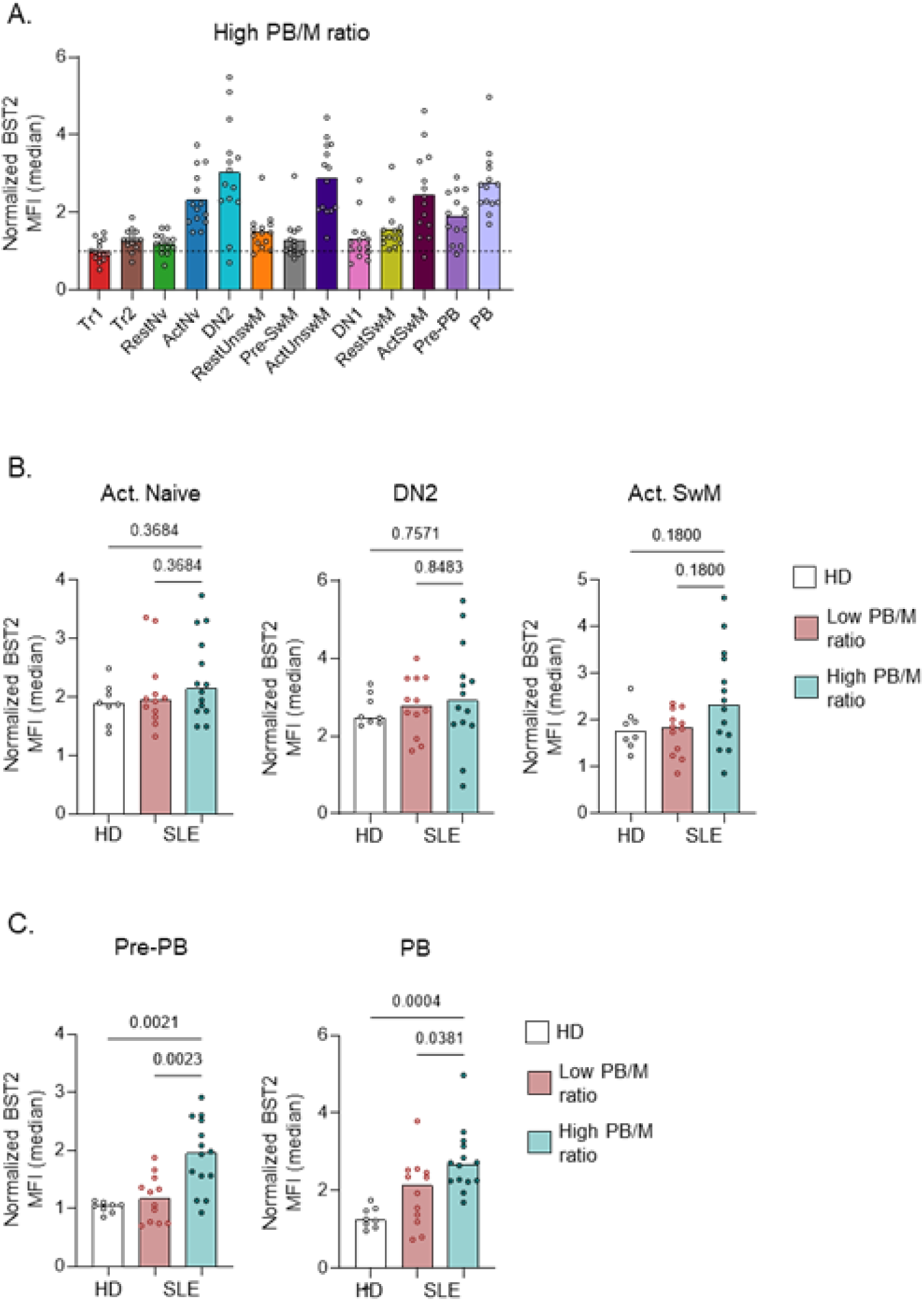
BST2 protein expression in B cell subsets. High dimensional spectral flow cytometry was used for analysis of BST2 expression within B cell subsets from SLE patients (n=26) and healthy donors (HD; n=8). Additional data on subset definitions is provided in Figure 2, S3 and S4. A) Normalized expression of BST2 in B cell subsets from patients with a high PB/M ratio. B,C) Comparison of normalized BST2 expression between patient groups and HD in subsets with high BST2 expression. Each dot indicates an individual, and the bars represent the median. P values were calculated using Kruskal- Wallis with FDR posthoc test (B,C).

**Figure S9:**
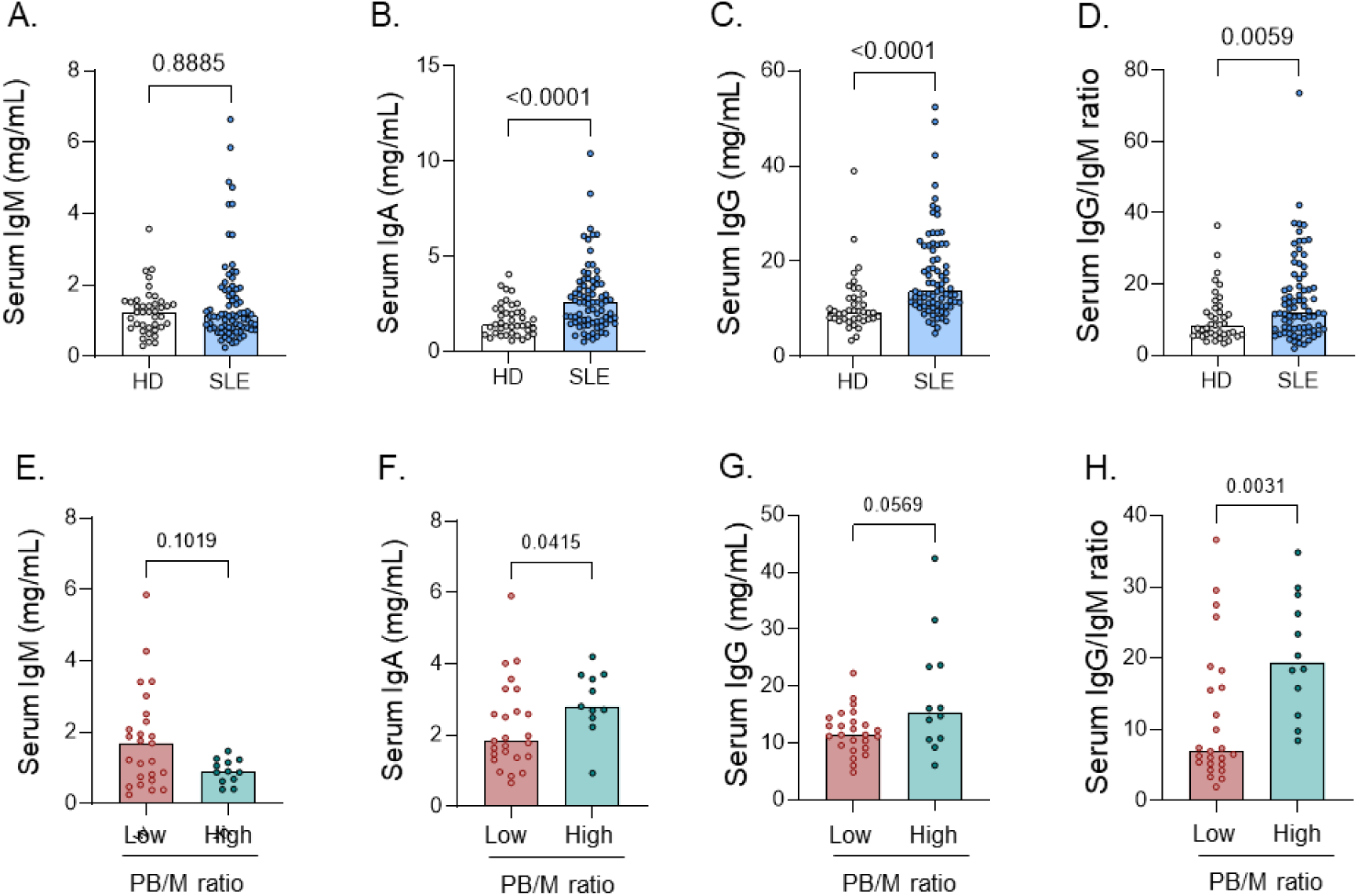
IFNAR expression by isotype and V gene distribution. BCR repertoire was determined using results from scRNA-seq (Figure 3) from 9 SLE patients, in 2 independent experiments. A) scRNAseq expression level of IFNAR1 and IFNAR2 in cells from each IGH isotype/subclass, separated by patient group. IGHD, IGHG3, and IGHG4 cells were not analysed due to low number of cells (Figure 4A). B) Expression of heavy chain V genes, expressed as percentage of total PB.

**Figure S8:**
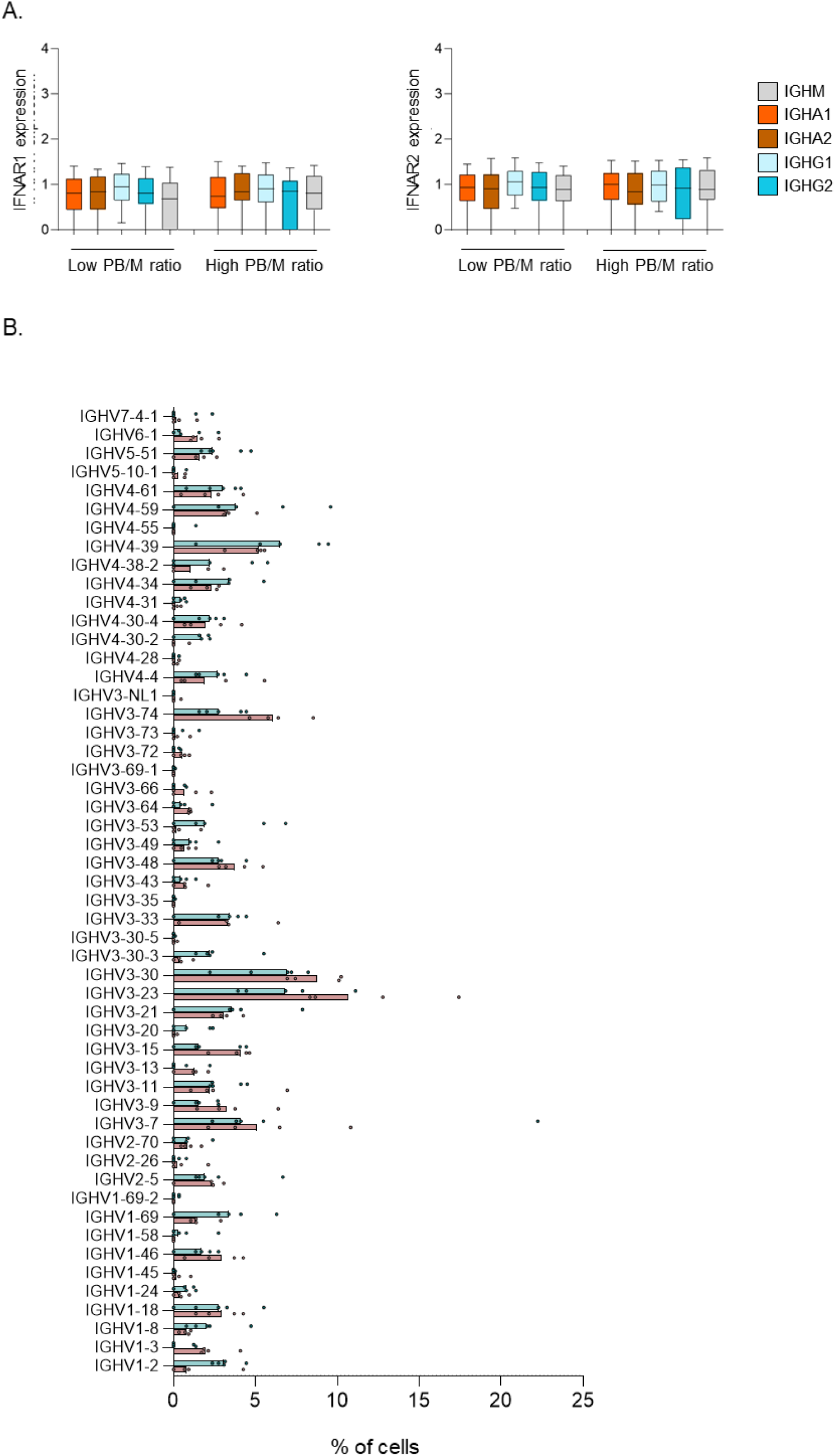
B cell hyperactivity results in hypergammaglobulinemia (replication cohort). Serum antibodies were measured in SLE patients (n=77) and healthy donors (n=40) using ELISA. SLE patients were split into two groups based on PB phenotype shown in Figure 1. A-H) Serum levels of total IgG, IgA, and IgM, and IgG:IgM ratio in SLE patients and healthy donors (A-D) and SLE patients split according to PB/CD27 group (E-H). Each dot indicates an individual, and the bars represent the median (A-H,J). P values were obtained using Mann- Whitney test (A-H).

**Figure S10:**
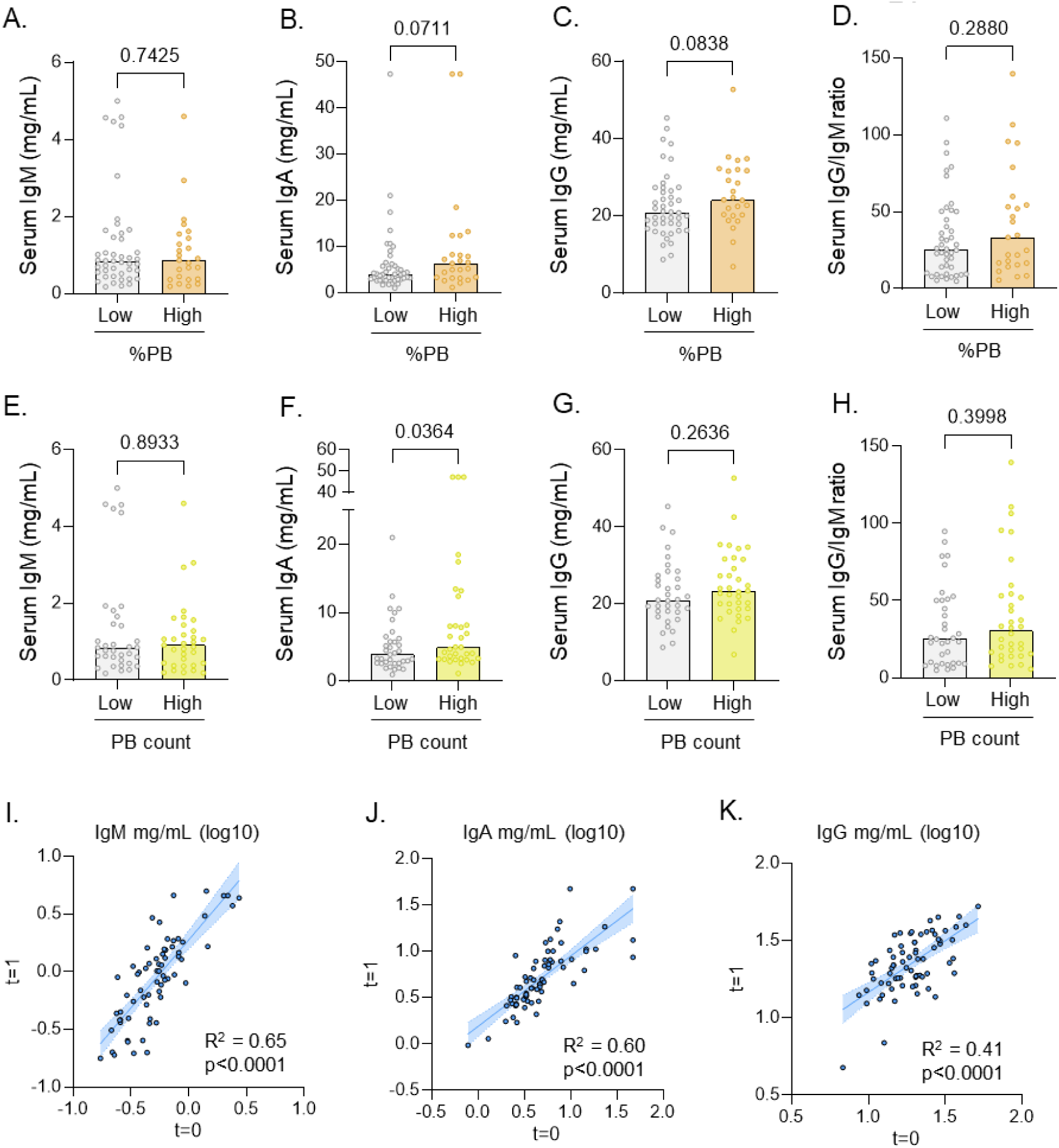
Hypergammaglobulinemia correlates less with PB count and %PB. Serum antibodies were measured in SLE patients (n=69) and healthy donors (n=40) using ELISA. SLE patients were split into two groups based on %PB and PB count. A-H) Serum levels of total IgG, IgA, and IgM, and IgG:IgM ratio in SLE patients split according to %PB (A-D) or PB count (E-H). I-K) Correlation of serum immunoglobulin levels in SLE patients measured at two timepoints ∼1 year apart (n=72). Each dot indicates an individual, and the bars represent the median (A-H). P values were obtained using Mann- Whitney test (A-H), or simple linear regression (I-K).

**Figure S11:**
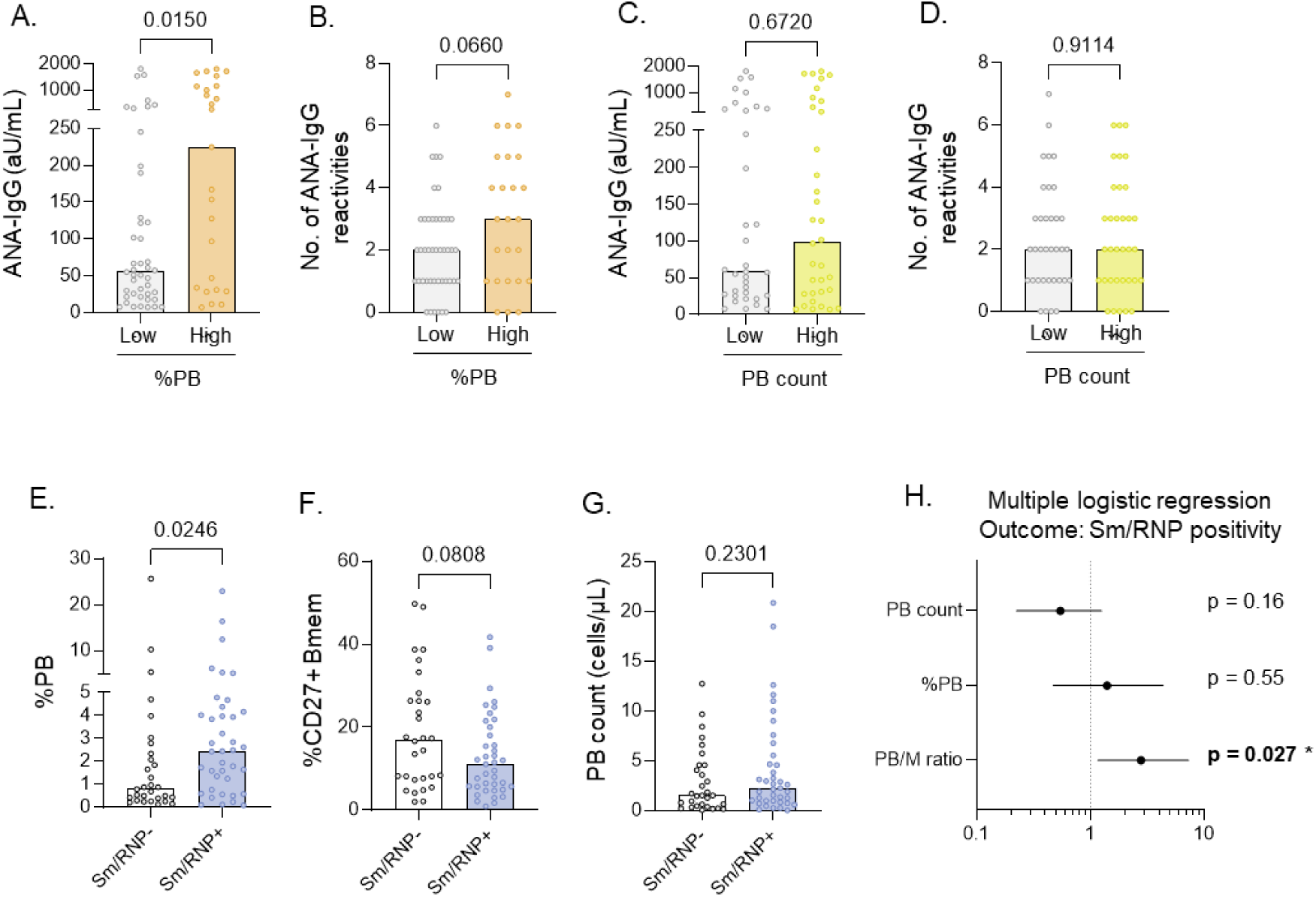
Sm/RNP reactivity correlates less with PB count and %PB. Serum autoantibodies were measured in SLE patients (n=69) using ELISA. Healthy donors (n=40) were used as negative controls. Autoantibody reactivities were categorized into three main antigen groups: chromatin (dsDNA, nucleosomes, histones); Sm/RNP (Sm, RNP70, U1-RNP complex), SS-A/B (SS-A, SS-B). SLE patients were split into two groups based on PB phenotype shown in Figure 1. A-D) Level of ANA-IgG and number of reactivities in SLE patients split according to %PB (A,B) and PB count (C,D). E-G) %PB, %Bmem, and PB count in SLE patients split according to Sm/RNP-IgG reactivity. H) Forest plot of multiple logistic regression model with Sm/RNP reactivity as outcome, and the Z-score of PB count, %PB, PB/M ratio, and cohort as predicting variables. Each dot indicates an individual, and the bars represent the median (A-G). P values were obtained using Mann- Whitney test (A-G), or multiple logistic regression (H).

**Figure S12:**
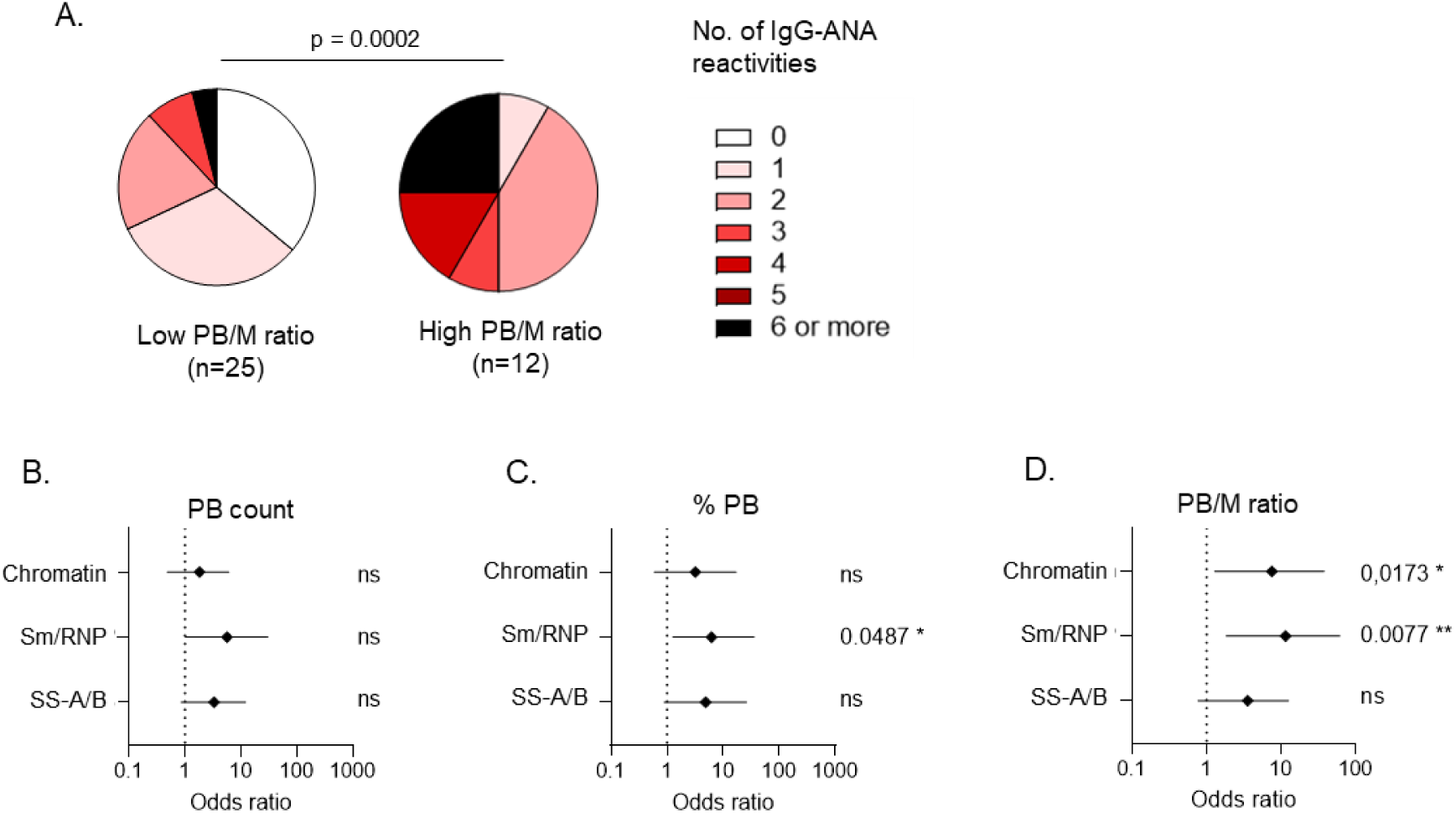
B cell hyperactivity is specifically associated with the presence of Sm/RNP autoantibodies (replication cohort). Serum autoantibodies were measured in SLE patients (n=37) using ELISA. Healthy donors (n=40) were used as negative controls. Autoantibody reactivities were categorized into three main antigen groups: chromatin (dsDNA, nucleosomes, histones); Sm/RNP (Sm, RNP70, U1-RNP complex), SS-A/B (SS-A, SS-B). SLE patients were split into two groups based on PB phenotype shown in Figure 1. A) Number of IgG-ANA reactivities in SLE patient groups. B-D) Forest plots of odds ratios with 95% confidence intervals obtained with Fisher’s exact test for specific ANA reactivity with the indicated PB groups. Details are shown in Table S7. P values were obtained using Mann-Whitney test (A), or Fisher’s exact test (B-D).

**Figure S13:**
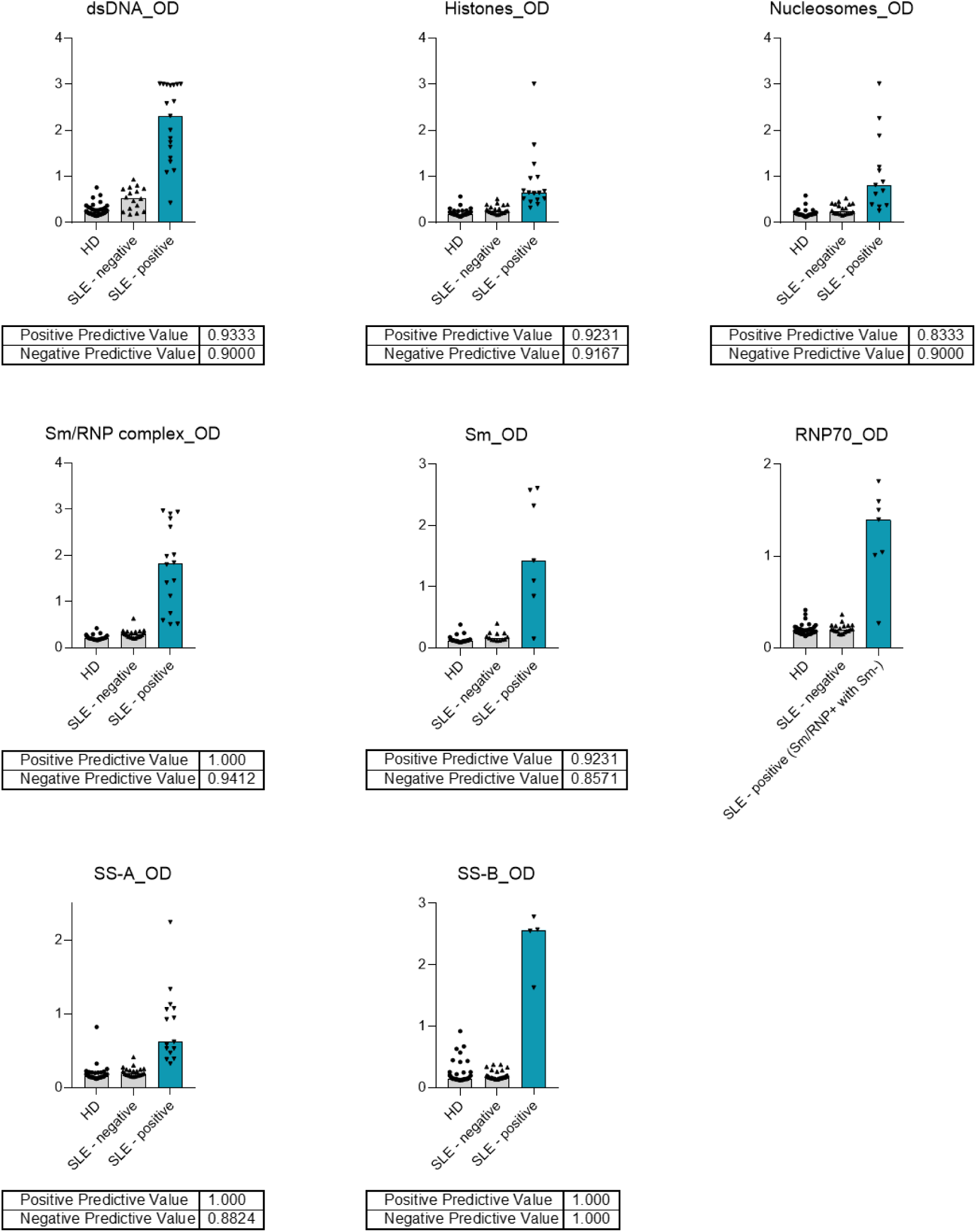
Validation of in-house ANA ELISAs. SLE patients negative or positive for the respective antigen were selected based on commercial ELISA or clinical diagnostic assays (Sm). Forty healthy donors were used to calculate a cutoff (mean + 4x SD) after which positive and negative predictive values were calculated using Fisher’s exact test. For RNP70, positive predictive value could not be calculated as the exact status of RNP70 reactivity was unknown (samples included Sm-RNP+ samples that were Sm-negative).

**Table S1:** Patient characteristics in PB/M low and PB/M high groups in Cohort 1 (Feinstein) The characteristics shown here are obtained during their initial assessment. All parameters are shown as the median and range or number and frequency within the patient group. None of the patients had used cytoxan or rituximab in the preceding 12 months. Statistical testing was done using Mann Whitney U test for linear parameters, chi-square test for categorical parameters with more than 2 categories (ethnicity) and Fisher’s exact test for categorical parameters with 2 categories (male/female and medication). ^1^ Immunosuppressants include Mycophenolate mofetil, Azathioprine and Methotrexate. PDN: Prednisone, HCQ: Hydroxychloroquine.

|  | Low PB/M ratio<br>(n=32) | High PB/M ratio<br>(n=40) | p value |
| --- | --- | --- | --- |
| Age (yrs) | 41 (21-64) | 41 (19-70) | 0.6745 |
| Male/female (% female) | 1/31 (97%) | 7/33 (83%) | 0.0684 |
| Ethnicity (self-identified) |  |  |  |
| African-American | 12 (38%) | 25 (63%) | 0.0570 |
| Hispanic | 13 (41%) | 9 (23%) | 0.1251 |
| Caucasian | 3 (9.4%) | 5 (13%) | 0.7254 |
| Asian | 4 (13%) | 0 (0%) | 0.0350 |
| Other | 0 (0%) | 1 (2.5%) | >0.9999 |
| Disease duration (yrs) | 10 (1-21) | 10 (0-36) | 0.4822 |
| Age at diagnosis (yrs) | 30 (9-56) | 25 (7-66) | 0.1666 |
| Current medication |  |  |  |
| PDN | 15 (47%) | 21 (53%) | 0.8128 |
| HCQ | 28 (88%) | 31 (78%) | 0.3611 |
| Immunosuppressants<sup>1</sup> | 14 (44%) | 19 (48%) | 0.8145 |

**Table S2:** Patient characteristics in low PB/M and high PB/M groups in Cohort 2 (Leiden) The characteristics shown here are obtained during their initial assessment. All parameters are shown as the median and range or number and frequency within the patient group. None of the patients had used cytoxan or rituximab in the preceding 12 months. Statistical testing was done using Mann Whitney U test for linear parameters, and Fisher’s exact test for categorical parameters with 2 categories (male/female and medication). ^1^ Immunosuppressants include Mycophenolate mofetil, Azathioprine and Methotrexate. PDN: Prednisone, HCQ: Hydroxychloroquine.

|  | Low PB/M ratio<br>(n=25) | High PB/M ratio<br>(n=12) | p value |
| --- | --- | --- | --- |
| Age (yrs) | 50 (22-76) | 47 (25-76) | 0.8792 |
| Male/female (% female) | 1/24 (96%) | 1/11 (92%) | >0.9999 |
| Disease duration (yrs) | 18 (2-39) | 21 (4-41) | 0.3550 |
| Age at diagnosis (yrs) | 31 (11-65) | 30 (13-40) | 0.6713 |
| Current medication |  |  |  |
| PDN | 3 (12%) | 6 (50%) | 0.0355 |
| HCQ | 19 (76%) | 10 (83%) | >0.9999 |
| Immunosuppressants<sup>1</sup> | 6 (24%) | 6 (50%) | 0.1138 |

**Table S3:** Disease activity in SLE patients from low PB/M and high PB/M groups. P value was obtained using Chi-square test.

|  | Low PB/M ratio<br>(n=32) | High PB/M ratio<br>(n=40) | p value |
| --- | --- | --- | --- |
| Inactive (SLEDAI-2k = 0) | 12 (38%) | 5 (13%) | **0.0293 \*** |
| Low (SLEDAI-2k = 1-4) | 14 (44%) | 19 (48%) |  |
| Moderate (SLEDAI-2k = 5-8) | 5 (16%) | 8 (20%) |  |
| High (SLEDAI-2k >8) | 1 (3%) | 8 (20%) |  |

**Table S4:** Clinical symptoms in SLE patients from low PB/M and high PB/M groups. P value was obtained using Fishers exact test.

|  | Low PB/M<br>ratio (n=32) | High PB/M<br>ratio (n=40) | p value |
| --- | --- | --- | --- |
| Renal | 2 (6%) | 7 (18%) | 0.2821 |
| Mucocutaneous | 7 (22%) | 14 (35%) | 0.2988 |
| Musculoskeletal | 3 (9%) | 7 (18%) | 0.4955 |
| Hematological | 0 (0%) | 6 (15%) | **0.0304 \*** |
| Other | 0 (0%) | 4 (10%) | 0.1216 |

**Table S5:**
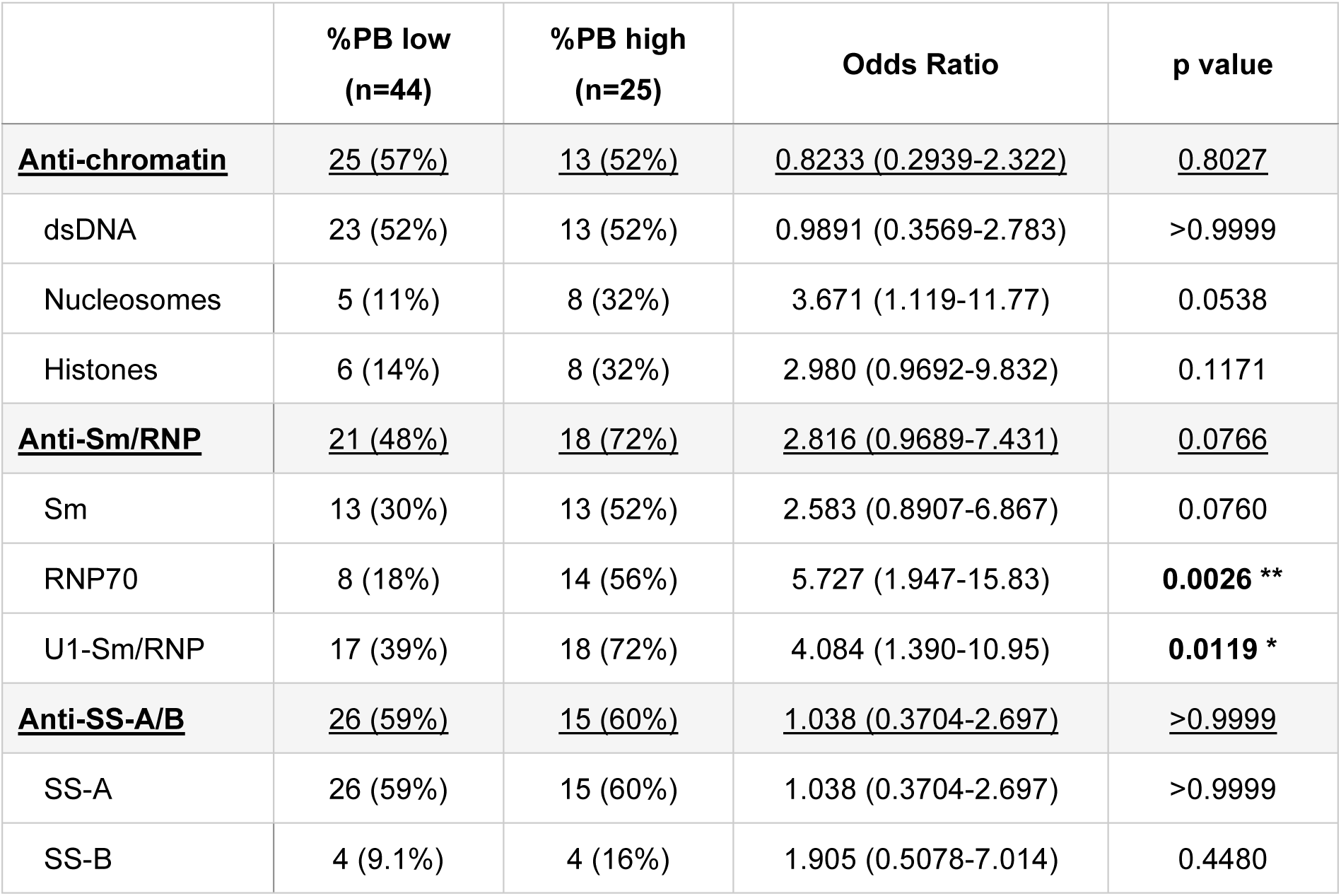
Frequency of ANA reactivities in SLE patients with a low and high %PB. Positivity of ANA-IgG for each indicated antigen was determined using ELISA. Anti-chromatin refers to all patients positive for 1 or more of the antigens dsDNA, Nucleosomes, and histones. Anti-Sm/RNP refers to all patients positive for 1 or more of the antigens Sm, and RNP70. Anti-SS-A/B refers to all patients positive for 1 or more of the antigens SS-A, and SS-B (all patients with SS-B were also positive for SS-A). P values were obtained using Fisher’s exact test.

|  | **%PB low**<br>**(n=44)** | **%PB high**<br>**(n=25)** | **Odds Ratio** | **p value** |
| --- | --- | --- | --- | --- |
| **<u>Anti-chromatin</u>** | **<u>25 (57%)</u>** | **<u>13 (52%)</u>** | **<u>0.8233 (0.2939-2.322)</u>** | **<u>0.8027</u>** |
| dsDNA | 23 (52%) | 13 (52%) | 0.9891 (0.3569-2.783) | >0.9999 |
| Nucleosomes | 5 (11%) | 8 (32%) | 3.671 (1.119-11.77) | 0.0538 |
| Histones | 6 (14%) | 8 (32%) | 2.980 (0.9692-9.832) | 0.1171 |
| **<u>Anti-Sm/RNP</u>** | **<u>21 (48%)</u>** | **<u>18 (72%)</u>** | **<u>2.816 (0.9689-7.431)</u>** | **<u>0.0766</u>** |
| Sm | 13 (30%) | 13 (52%) | 2.583 (0.8907-6.867) | 0.0760 |
| RNP70 | 8 (18%) | 14 (56%) | 5.727 (1.947-15.83) | **0.0026 \*\*** |
| U1-Sm/RNP | 17 (39%) | 18 (72%) | 4.084 (1.390-10.95) | **0.0119 \*** |
| **<u>Anti-SS-A/B</u>** | **<u>26 (59%)</u>** | **<u>15 (60%)</u>** | **<u>1.038 (0.3704-2.697)</u>** | **<u>>0.9999</u>** |
| SS-A | 26 (59%) | 15 (60%) | 1.038 (0.3704-2.697) | >0.9999 |
| SS-B | 4 (9.1%) | 4 (16%) | 1.905 (0.5078-7.014) | 0.4480 |

**Table S6:** Frequency of ANA reactivities in SLE patients with a low and high PB count. Positivity of ANA-IgG for each indicated antigen was determined using ELISA. Anti-chromatin refers to all patients positive for 1 or more of the antigens dsDNA, Nucleosomes, and histones. Anti-Sm/RNP refers to all patients positive for 1 or more of the antigens Sm, and RNP70. Anti-SS-A/B refers to all patients positive for 1 or more of the antigens SS-A, and SS-B (all patients with SS-B were also positive for SS-A). P values were obtained using Fisher’s exact test.

|  | **PB count low**<br>**(n=35)** | **PB count**<br>**high (n=34)** | **Odds Ratio** | **p value** |
| --- | --- | --- | --- | --- |
| **<u>Anti-chromatin</u>** | **<u>22 (63%)</u>** | **<u>16 (47%)</u>** | **<u>0.5253 (0.1915-1.345)</u>** | **<u>0.2301</u>** |
| dsDNA | 21 (60%) | 15 (44%) | 0.5263 (0.1943-1.337) | 0.2316 |
| Nucleosomes | 5 (14%) | 8 (23%) | 1.778 (0.5649-5.531) | 0.5401 |
| Histones | 7 (20%) | 7 (21%) | 1.037 (0.3340-3.220) | >0.9999 |
| **<u>Anti-Sm/RNP</u>** | **<u>17 (49%)</u>** | **<u>22 (65%)</u>** | **<u>1.941 (0.7524-4.925)</u>** | **<u>0.2270</u>** |
| Sm | 11 (31%) | 15 (44%) | 1.722 (0.6402-4.295) | 0.3261 |
| RNP70 | 7 (20%) | 15 (44%) | 3.158 (1.043-8.526) | **0.0406 \*** |
| U1-Sm/RNP | 15 (43%) | 20 (59%) | 1.905 (0.7510-5.157) | 0.2316 |
| **<u>Anti-SS-A/B</u>** | **<u>23 (66%)</u>** | **<u>18 (53%)</u>** | **<u>0.5870 (0.2271-1.537)</u>** | **<u>0.3319</u>** |
| SS-A | 23 (66%) | 18 (53%) | 0.5870 (0.2271-1.537) | 0.3319 |
| SS-B | 5 (14%) | 3 (8.8%) | 0.5806 (0.1448-2.546) | 0.7096 |

**Table S7:** Frequency of ANA reactivities in SLE patients from low PB/M and high PB/M groups in the replication cohort (Leiden). Positivity of ANA-IgG for each indicated antigen was determined using ELISA. Anti-chromatin refers to all patients positive for 1 or more of the antigens dsDNA, Nucleosomes, and histones. Anti-Sm/RNP refers to all patients positive for 1 or more of the antigens Sm, and RNP70. Anti-SS-A/B refers to all patients positive for 1 or more of the antigens SS-A, and SS-B (all patients with SS-B were also positive for SS-A). P values were obtained using Fisher’s exact test.

|  | Low PB/M<br>ratio (n=25) | High PB/M<br>ratio (n=12) | Odds Ratio | p value |
| --- | --- | --- | --- | --- |
| **<u>Anti-chromatin</u>** | <u>10 (40%)</u> | <u>10 (83%)</u> | <u>7.50 (1.29-37.78)</u> | **<u>0.0173 \*</u>** |
| dsDNA | 8 (32%) | 10 (83%) | 10.63 (1.77-53.42) | **0.0051 \*\*** |
| Nucleosomes | 3 (12%) | 3 (25%) | 2.44 (0.48-11.76) | 0.3666 |
| Histones | 4 (16%) | 5 (42%) | 3.75 (0.73-14.61) | 0.1161 |
| **<u>Anti-Sm/RNP</u>** | <u>2 (8%)</u> | <u>6 (50%)</u> | <u>11.50 (1.84-61.50)</u> | **<u>0.0077 \*\*</u>** |
| Sm | 2 (8%) | 6 (50%) | 11.50 (1.84-61.50) | **0.0077 \*\*** |
| RNP70 | 0 (0%) | 2 (17%) | Inf (1.01-inf) | 0.0991 |
| U1-Sm/RNP | 1 (4%) | 6 (50%) | 24.00 (2.47-283.0) | **0.0023 \*\*** |
| **<u>Anti-SS-A/B</u>** | <u>9 (36%)</u> | <u>8 (67%)</u> | <u>3.56 (0.77-12.57)</u> | <u>0.1575</u> |
| SS-A | 9 (36%) | 8 (67%) | 3.56 (0.77-12.57) | 0.1575 |
| SS-B | 4 (16%) | 6 (50%) | 5.25 (1.16-20.51) | **0.0486 \*** |

**Table S8:**
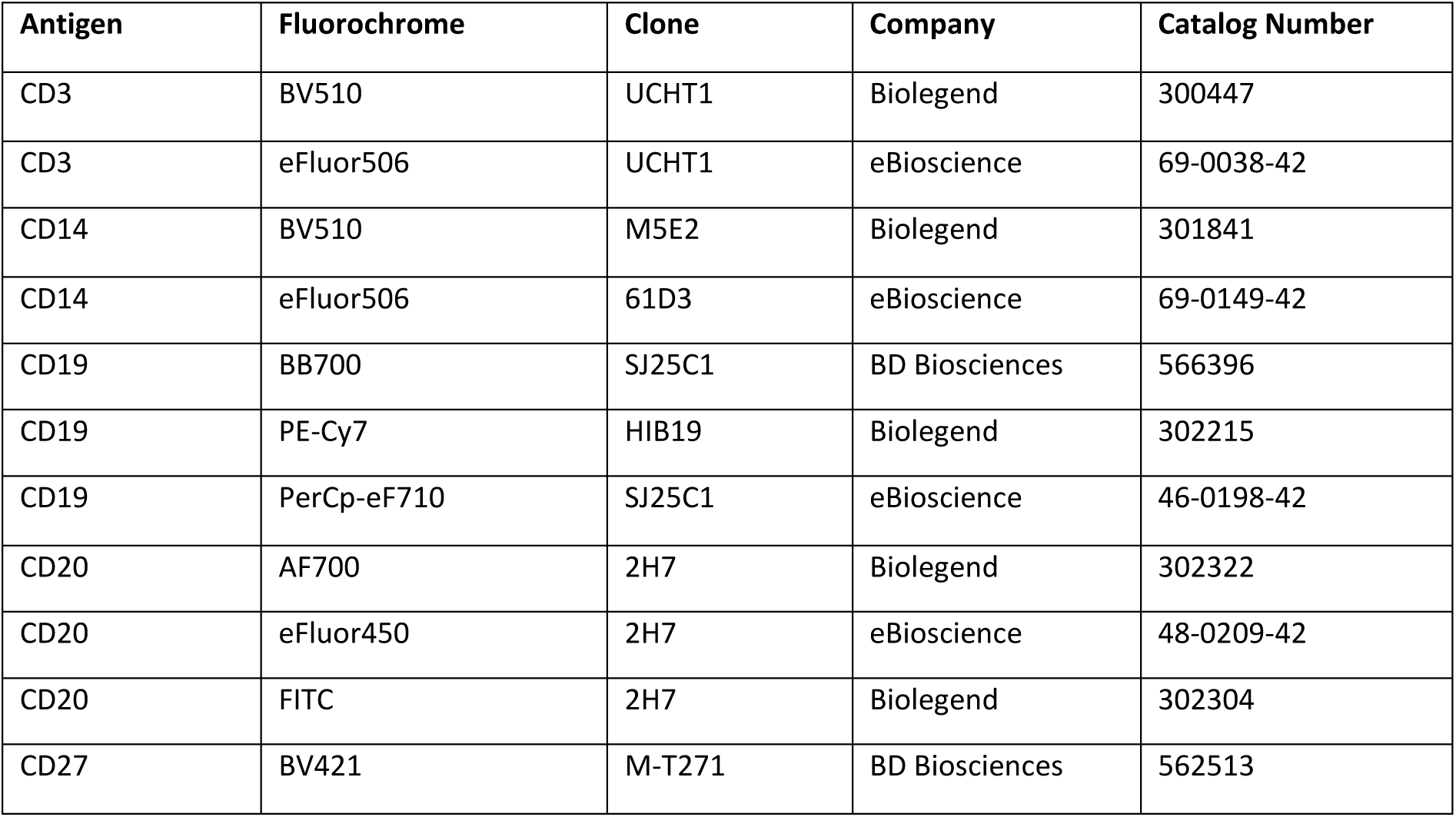
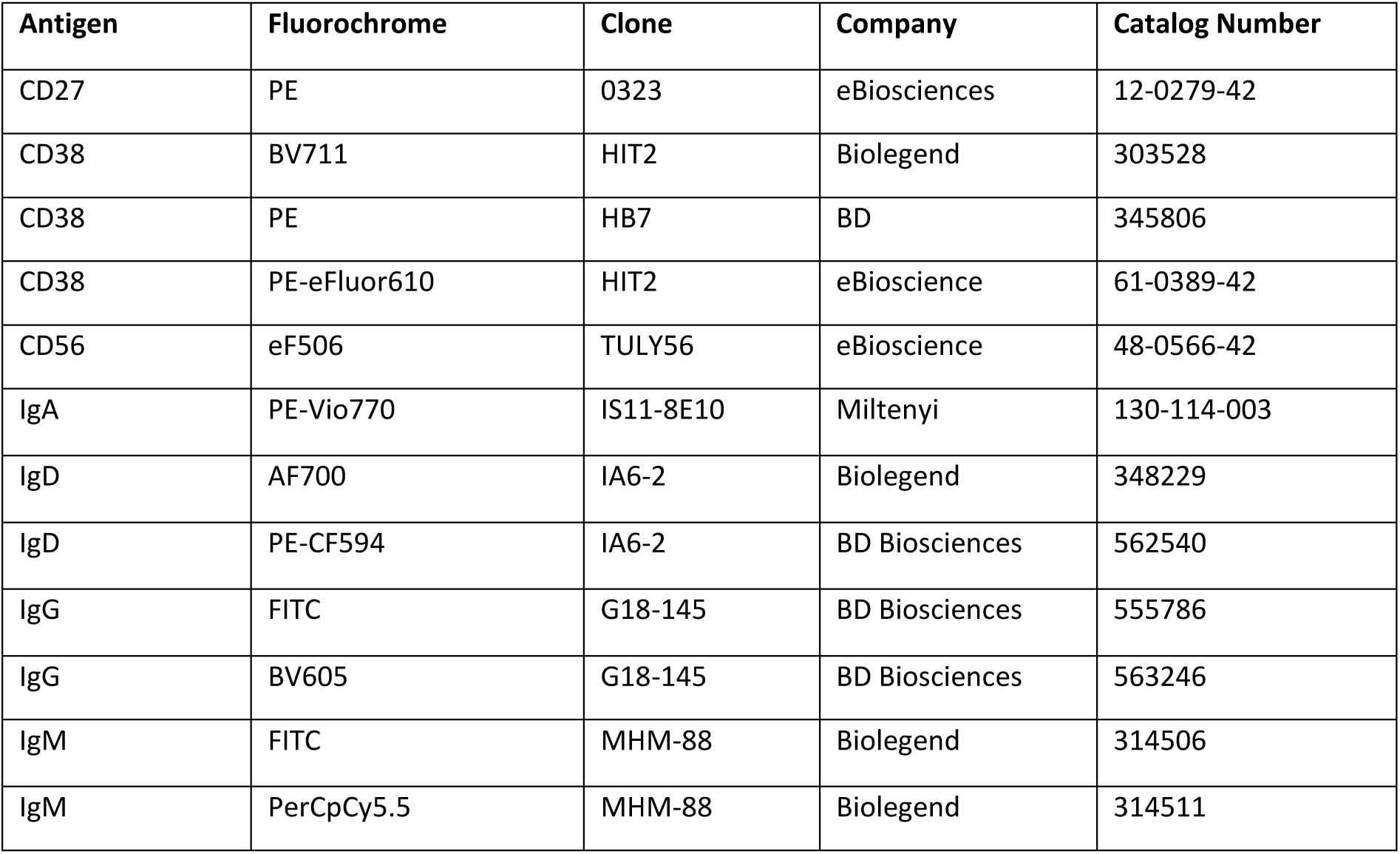
Conventional flow cytometry & FACS antibodies

**Table S9:** Spectral flow cytometry antibodies

| Antigen | Fluorochrome | Clone | Company | Catalog Number |
| --- | --- | --- | --- | --- |
| CD1c (BDCA1) | PE | AD5-8E7 | Miltenyi | 120-000-889 |
| CD1d | PE-Cy7 | 51.1 | Biolegend | 350309 |
| CD3 | eFluor 506 | UCHT1 | eBioscience | 69-0038-42 |
| CD11c | FITC | 3.9 | Biolegend | 301604 |
| CD14 | eFluor 506 | 61D3 | eBioscience | 69-0149-42 |
| CD19 | BV570 | HIB19 | Biolegend | 302236 |
| CD20 | BUV395 | 2H7 | BD | 563782 |
| CD21 | BUV805 | B-ly4 | BD | 742008 |
| CD24 | BUV496 | ML5 | BD | 741143 |
| CD27 | APC-Fire810 | QA17A18 | Biolegend | 393214 |
| CD32B/C | AF647 | 4F5 | Biolegend | Custom labeled by Biolegend |
| CD38 | BUV563 | HB7 | BD | 741446 |
| CD40 | PerCpCy5.5 | 5C3 | Biolegend | 334315 |
| CD45RB | BUV615 | MT4 (6B6) | BD | 751482 |
| CD49D | BV711 | 9F10 | Biolegend | 304331 |
| CD69 | PE-Fire640 | FN50 | Biolegend | 310959 |
| CD70 | PE | Ki-24 | BD Biosciences | 555835 |
| CD80 | APC-R700 | L307.4 | BD | 565157 |
| CD86 | BV650 | IT2.2 | Biolegend | 305427 |
| CD138 | BV711 | MI15 | Biolegend | 356521 |
| CD183 (CXCR3) | AF488 | G025H7 | Biolegend | 353709 |
| CD183 (CXCR3) | BV785 | G025H7 | Biolegend | 353737 |
| CD197 (CCR7) | APC-Fire750 | G043H7 | Biolegend | 353245 |
| CD317 (BST2) | PE-Cy7 | RS38E | Biolegend | 348415 |
| CXCR5 | BV750 | J252D4 | Biolegend | 356941 |
| HLA-DR | PE-Fire810 | L243 | Biolegend | 307683 |
| IgD | BV480 | IA6-2 | BD | 566138 |
| IgM | SuperBright436 | SA-DA4 | eBioscience | 62-9998-42 |
| IgG | BV421 | G18-145 | BD | 562581 |
| IgA | VioBlue | IS11-8E10 | Miltenyi | 130-113-479 |
| Ki67 | PerCp-eF70 | 20Raj1 | eBioscience | 46-5699-42 |
| S1PR4 | AF488 | 1012512 | R&D systems | FAB10321G |

**Table S10:**
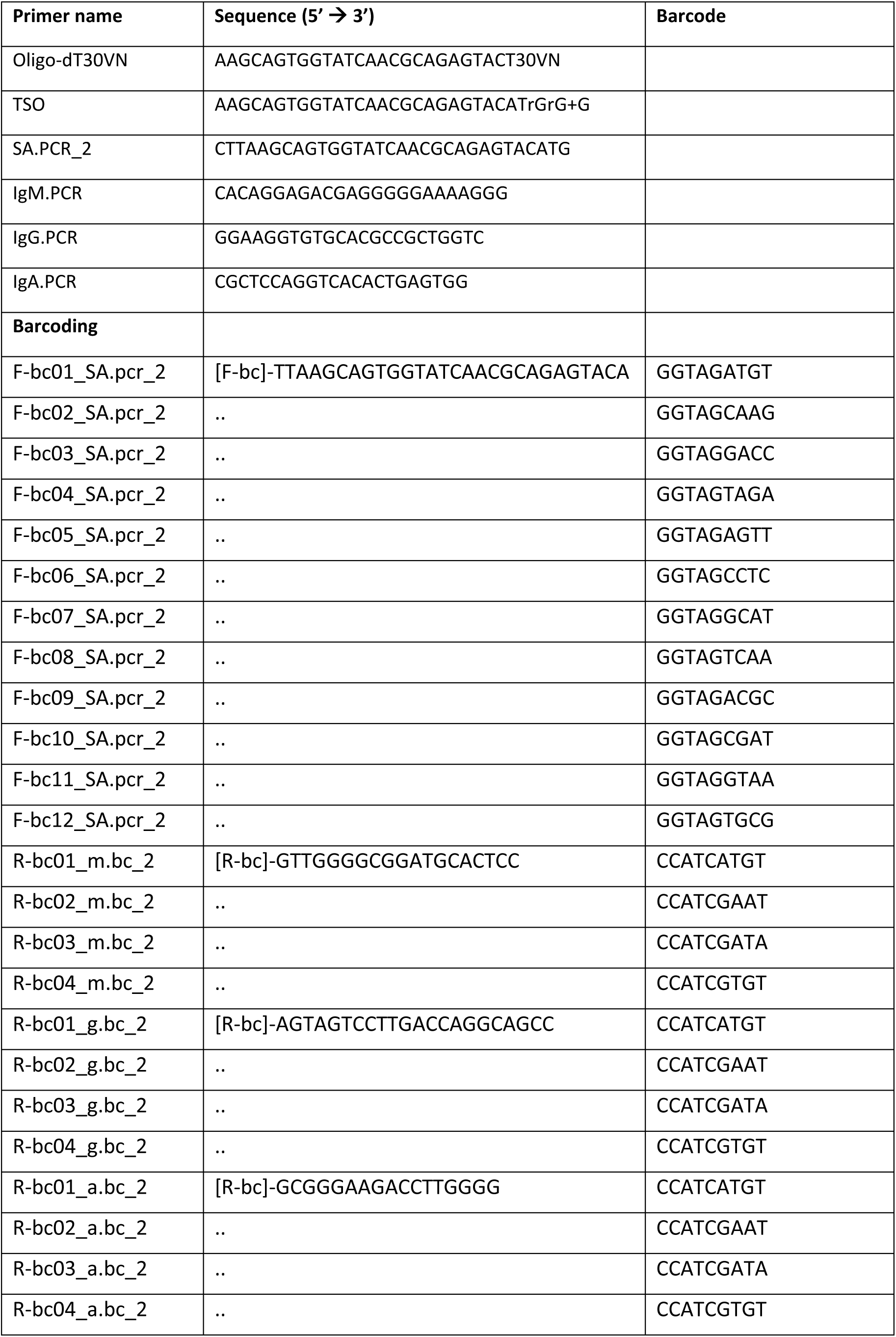
Primer sequences for 5’ RACE PCR

## REFERENCES

1. Yu C, Gershwin ME, Chang C. Diagnostic criteria for systemic lupus erythematosus: a critical review. J Autoimmun. 2014;48–49:10–3.

2. Suurmond J, Diamond B. Autoantibodies in systemic autoimmune diseases: specificity and pathogenicity. J Clin Invest. 2015;125(6):2194–202.

3. Aringer M, Costenbader K, Daikh D, Brinks R, Mosca M, Ramsey-Goldman R, et al. 2019 European League Against Rheumatism/American College of Rheumatology Classification Criteria for Systemic Lupus Erythematosus. Arthritis Rheumatol. 2019;71(9):1400–12.

4. Malkiel S, Barlev AN, Atisha-Fregoso Y, Suurmond J, Diamond B. Plasma Cell Differentiation Pathways in Systemic Lupus Erythematosus. Front Immunol. 2018;9:427.

5. ter Borg EJ, Horst G, Hummel EJ, Limburg PC, Kallenberg CG. Measurement of increases in anti-double-stranded DNA antibody levels as a predictor of disease exacerbation in systemic lupus erythematosus. A long-term, prospective study. Arthritis Rheum. 1990;33(5):634–43.

6. Pisetsky DS, Lipsky PE. New insights into the role of antinuclear antibodies in systemic lupus erythematosus. Nat Rev Rheumatol. 2020;16(10):565–79.

7. Muller F, Taubmann J, Bucci L, Wilhelm A, Bergmann C, Volkl S, et al. CD19 CAR T-Cell Therapy in Autoimmune Disease - A Case Series with Follow-up. N Engl J Med. 2024;390(8):687–700.

8. Suurmond J, Atisha-Fregoso Y, Marasco E, Barlev AN, Ahmed N, Calderon SA, et al. Loss of an IgG plasma cell checkpoint in patients with lupus. J Allergy Clin Immunol. 2019;143(4):1586–97.

9. Dorner T, Giesecke C, Lipsky PE. Mechanisms of B cell autoimmunity in SLE. Arthritis Res Ther. 2011;13(5):243.

10. Ye Z, Jiang Y, Sun D, Zhong W, Zhao L, Jiang Z. The Plasma Interleukin (IL)-35 Level and Frequency of Circulating IL-35(+) Regulatory B Cells are Decreased in a Cohort of Chinese Patients with New-onset Systemic Lupus Erythematosus. Sci Rep. 2019;9(1):13210.

11. Banchereau R, Hong S, Cantarel B, Baldwin N, Baisch J, Edens M, et al. Personalized Immunomonitoring Uncovers Molecular Networks that Stratify Lupus Patients. Cell. 2016;165(6):1548–50.

12. Odendahl M, Jacobi A, Hansen A, Feist E, Hiepe F, Burmester GR, et al. Disturbed peripheral B lymphocyte homeostasis in systemic lupus erythematosus. J Immunol. 2000;165(10):5970–9.

13. Arce E, Jackson DG, Gill MA, Bennett LB, Banchereau J, Pascual V. Increased frequency of pre-germinal center B cells and plasma cell precursors in the blood of children with systemic lupus erythematosus. J Immunol. 2001;167(4):2361–9.

14. Jacobi AM, Odendahl M, Reiter K, Bruns A, Burmester GR, Radbruch A, et al. Correlation between circulating CD27high plasma cells and disease activity in patients with systemic lupus erythematosus. Arthritis Rheum. 2003;48(5):1332–42.

15. Klinman DM, Shirai A, Ishigatsubo Y, Conover J, Steinberg AD. Quantitation of IgM- and IgG- secreting B cells in the peripheral blood of patients with systemic lupus erythematosus. Arthritis Rheum. 1991;34(11):1404–10.

16. Barlev AN, Malkiel S, Kurata-Sato I, Dorjee AL, Suurmond J, Diamond B. FcgammaRIIB regulates autoantibody responses by limiting marginal zone B cell activation. J Clin Invest. 2022;132(17).

17. Samuelson EM, Laird RM, Maue AC, Rochford R, Hayes SM. Blk haploinsufficiency impairs the development, but enhances the functional responses, of MZ B cells. Immunol Cell Biol. 2012;90(6):620–9.

18. He Y, Gallman AE, Xie C, Shen Q, Ma J, Wolfreys FD, et al. P2RY8 variants in lupus patients uncover a role for the receptor in immunological tolerance. J Exp Med. 2022;219(1).

19. Brown GJ, Canete PF, Wang H, Medhavy A, Bones J, Roco JA, et al. TLR7 gain-of-function genetic variation causes human lupus. Nature. 2022;605(7909):349–56.

20. Suurmond J, Atisha-Fregoso Y, Barlev AN, Calderon SA, Mackay MC, Aranow C, et al. Patterns of ANA+ B cells for SLE patient stratification. JCI Insight. 2019;4(9).

21. Tipton CM, Fucile CF, Darce J, Chida A, Ichikawa T, Gregoretti I, et al. Diversity, cellular origin and autoreactivity of antibody-secreting cell population expansions in acute systemic lupus erythematosus. Nat Immunol. 2015;16(7):755–65.

22. Wehr C, Eibel H, Masilamani M, Illges H, Schlesier M, Peter HH, et al. A new CD21low B cell population in the peripheral blood of patients with SLE. Clin Immunol. 2004;113(2):161–71.

23. Jenks SA, Cashman KS, Zumaquero E, Marigorta UM, Patel AV, Wang X, et al. Distinct Effector B Cells Induced by Unregulated Toll-like Receptor 7 Contribute to Pathogenic Responses in Systemic Lupus Erythematosus. Immunity. 2018;49(4):725–39 e6.

24. Dieudonne Y, Gies V, Guffroy A, Keime C, Bird AK, Liesveld J, et al. Transitional B cells in quiescent SLE: An early checkpoint imprinted by IFN. J Autoimmun. 2019;102:150–8.

25. Sanz I, Wei C, Jenks SA, Cashman KS, Tipton C, Woodruff MC, et al. Challenges and Opportunities for Consistent Classification of Human B Cell and Plasma Cell Populations. Front Immunol. 2019;10:2458.

26. Siu JHY, Pitcher MJ, Tull TJ, Velounias RL, Guesdon W, Montorsi L, et al. Two subsets of human marginal zone B cells resolved by global analysis of lymphoid tissues and blood. Sci Immunol. 2022;7(69):eabm9060.

27. Priest DG, Ebihara T, Tulyeu J, Sondergaard JN, Sakakibara S, Sugihara F, et al. Atypical and non-classical CD45RB(lo) memory B cells are the majority of circulating SARS-CoV-2 specific B cells following mRNA vaccination or COVID-19. Nat Commun. 2024;15(1):6811.

28. Cooper GS, Parks CG, Treadwell EL, St Clair EW, Gilkeson GS, Cohen PL, et al. Differences by race, sex and age in the clinical and immunologic features of recently diagnosed systemic lupus erythematosus patients in the southeastern United States. Lupus. 2002;11(3):161–7.

29. Menard LC, Habte S, Gonsiorek W, Lee D, Banas D, Holloway DA, et al. B cells from African American lupus patients exhibit an activated phenotype. JCI Insight. 2016;1(9):e87310.

30. Navarra SV, Guzman RM, Gallacher AE, Hall S, Levy RA, Jimenez RE, et al. Efficacy and safety of belimumab in patients with active systemic lupus erythematosus: a randomised, placebo- controlled, phase 3 trial. Lancet. 2011;377(9767):721–31.

31. Saiki O, Saeki Y, Tanaka T, Doi S, Hara H, Negoro S, et al. Development of selective IgM deficiency in systemic lupus erythematosus patients with disease of long duration. Arthritis Rheum. 1987;30(11):1289–92.

32. Gronwall C, Hardt U, Gustafsson JT, Elvin K, Jensen-Urstad K, Kvarnstrom M, et al. Depressed serum IgM levels in SLE are restricted to defined subgroups. Clin Immunol. 2017;183:304–15.

33. Petri M, Fu W, Ranger A, Allaire N, Cullen P, Magder LS, et al. Association between changes in gene signatures expression and disease activity among patients with systemic lupus erythematosus. BMC Med Genomics. 2019;12(1):4.

34. Scharer CD, Blalock EL, Mi T, Barwick BG, Jenks SA, Deguchi T, et al. Epigenetic programming underpins B cell dysfunction in human SLE. Nat Immunol. 2019;20(8):1071–82.

35. Horisberger A, Humbel M, Fluder N, Bellanger F, Fenwick C, Ribi C, et al. Measurement of circulating CD21(-)CD27(-) B lymphocytes in SLE patients is associated with disease activity independently of conventional serological biomarkers. Sci Rep. 2022;12(1):9189.

36. Zurbuchen Y, Michler J, Taeschler P, Adamo S, Cervia C, Raeber ME, et al. Human memory B cells show plasticity and adopt multiple fates upon recall response to SARS-CoV-2. Nat Immunol. 2023;24(6):955–65.

37. Lau D, Lan LY, Andrews SF, Henry C, Rojas KT, Neu KE, et al. Low CD21 expression defines a population of recent germinal center graduates primed for plasma cell differentiation. Sci Immunol. 2017;2(7).

38. Chen W, Hong SH, Jenks SA, Anam FA, Tipton CM, Woodruff MC, et al. Distinct transcriptomes and autocrine cytokines underpin maturation and survival of antibody-secreting cells in systemic lupus erythematosus. Nat Commun. 2024;15(1):1899.

39. Dogan I, Bertocci B, Vilmont V, Delbos F, Megret J, Storck S, et al. Multiple layers of B cell memory with different effector functions. Nat Immunol. 2009;10(12):1292–9.

40. Mesin L, Schiepers A, Ersching J, Barbulescu A, Cavazzoni CB, Angelini A, et al. Restricted Clonality and Limited Germinal Center Reentry Characterize Memory B Cell Reactivation by Boosting. Cell. 2020;180(1):92–106 e11.

41. Douagi I, Gujer C, Sundling C, Adams WC, Smed-Sorensen A, Seder RA, et al. Human B cell responses to TLR ligands are differentially modulated by myeloid and plasmacytoid dendritic cells. J Immunol. 2009;182(4):1991–2001.

42. Domeier PP, Chodisetti SB, Schell SL, Kawasawa YI, Fasnacht MJ, Soni C, et al. B-Cell-Intrinsic Type 1 Interferon Signaling Is Crucial for Loss of Tolerance and the Development of Autoreactive B Cells. Cell Rep. 2018;24(2):406–18.

43. Lovgren T, Eloranta ML, Kastner B, Wahren-Herlenius M, Alm GV, Ronnblom L. Induction of interferon-alpha by immune complexes or liposomes containing systemic lupus erythematosus autoantigen- and Sjogren’s syndrome autoantigen-associated RNA. Arthritis Rheum. 2006;54(6):1917–27.

44. Bave U, Alm GV, Ronnblom L. The combination of apoptotic U937 cells and lupus IgG is a potent IFN-alpha inducer. J Immunol. 2000;165(6):3519–26.

45. Hubbard EL, Pisetsky DS, Lipsky PE. Anti-RNP antibodies are associated with the interferon gene signature but not decreased complement levels in SLE. Ann Rheum Dis. 2022;81(5):632–43.

46. Jego G, Palucka AK, Blanck JP, Chalouni C, Pascual V, Banchereau J. Plasmacytoid dendritic cells induce plasma cell differentiation through type I interferon and interleukin 6. Immunity. 2003;19(2):225–34.

47. Kirou KA, Lee C, George S, Louca K, Peterson MG, Crow MK. Activation of the interferon- alpha pathway identifies a subgroup of systemic lupus erythematosus patients with distinct serologic features and active disease. Arthritis Rheum. 2005;52(5):1491–503.

48. Catalina MD, Bachali P, Yeo AE, Geraci NS, Petri MA, Grammer AC, et al. Patient ancestry significantly contributes to molecular heterogeneity of systemic lupus erythematosus. JCI Insight. 2020;5(15).

49. Bunce CM, Drayson MT. Dissecting racial disparities in multiple myeloma-clues from differential immunoglobulin levels. Blood Cancer J. 2020;10(4):44.

50. Alarcon GS, Calvo-Alen J, McGwin G, Jr., Uribe AG, Toloza SM, Roseman JM, et al. Systemic lupus erythematosus in a multiethnic cohort: LUMINA XXXV. Predictive factors of high disease activity over time. Ann Rheum Dis. 2006;65(9):1168–74.

51. Somers EC, Marder W, Cagnoli P, Lewis EE, DeGuire P, Gordon C, et al. Population-based incidence and prevalence of systemic lupus erythematosus: the Michigan Lupus Epidemiology and Surveillance program. Arthritis Rheumatol. 2014;66(2):369–78.

52. Gladman DD, Ibanez D, Urowitz MB. Systemic lupus erythematosus disease activity index 2000. J Rheumatol. 2002;29(2):288–91.

53. Van Gassen S, Gaudilliere B, Angst MS, Saeys Y, Aghaeepour N. CytoNorm: A Normalization Algorithm for Cytometry Data. Cytometry A. 2020;97(3):268–78.

54. Hao Y, Hao S, Andersen-Nissen E, Mauck WM, 3rd, Zheng S, Butler A, et al. Integrated analysis of multimodal single-cell data. Cell. 2021;184(13):3573–87 e29.

55. Aran D, Looney AP, Liu L, Wu E, Fong V, Hsu A, et al. Reference-based analysis of lung single- cell sequencing reveals a transitional profibrotic macrophage. Nat Immunol. 2019;20(2):163–72.

56. Monaco G, Lee B, Xu W, Mustafah S, Hwang YY, Carre C, et al. RNA-Seq Signatures Normalized by mRNA Abundance Allow Absolute Deconvolution of Human Immune Cell Types. Cell Rep. 2019;26(6):1627–40 e7.

57. Stoeckius M, Zheng S, Houck-Loomis B, Hao S, Yeung BZ, Mauck WM, 3rd, et al. Cell Hashing with barcoded antibodies enables multiplexing and doublet detection for single cell genomics. Genome Biol. 2018;19(1):224.

58. Reshef YA, Rumker L, Kang JB, Nathan A, Korsunsky I, Asgari S, et al. Co-varying neighborhood analysis identifies cell populations associated with phenotypes of interest from single- cell transcriptomics. Nat Biotechnol. 2022;40(3):355–63.

59. Merico D, Isserlin R, Stueker O, Emili A, Bader GD. Enrichment map: a network-based method for gene-set enrichment visualization and interpretation. PLoS One. 2010;5(11):e13984.

60. Subramanian A, Tamayo P, Mootha VK, Mukherjee S, Ebert BL, Gillette MA, et al. Gene set enrichment analysis: a knowledge-based approach for interpreting genome-wide expression profiles. Proc Natl Acad Sci U S A. 2005;102(43):15545–50.

61. Catalina MD, Bachali P, Geraci NS, Grammer AC, Lipsky PE. Gene expression analysis delineates the potential roles of multiple interferons in systemic lupus erythematosus. Commun Biol. 2019;2:140.

62. Koning MT, Kielbasa SM, Boersma V, Buermans HPJ, van der Zeeuw SAJ, van Bergen CAM, et al. ARTISAN PCR: rapid identification of full-length immunoglobulin rearrangements without primer binding bias. Br J Haematol. 2017;178(6):983–6.

63. Slot LM, Vergroesen RD, Kerkman PF, Staudinger E, Reijm S, van Dooren HJ, et al. Light chain skewing in autoantibodies and B-cell receptors of the citrullinated antigen-binding B-cell response in rheumatoid arthritis. PLoS One. 2021;16(3):e0247847.

64. Song L, Cohen D, Ouyang Z, Cao Y, Hu X, Liu XS. TRUST4: immune repertoire reconstruction from bulk and single-cell RNA-seq data. Nat Methods. 2021;18(6):627–30.

